# Identification of candidate biomarkers and therapeutic agents for heart failure by bioinformatics analysis

**DOI:** 10.1101/2021.01.24.428028

**Authors:** Basavaraj Vastrad, Anandkumar Tengli, Chanabasayya Vastrad

## Abstract

Heart failure (HF) is a heterogeneous clinical syndrome and affects millions of people all over the world. HF occurs when the cardiac overload and injury, which is a worldwide complaint. The aim of this study was to screen and verify hub genes involved in developmental HF as well as to explore active drug molecules. The expression profiling by high throughput sequencing of GSE141910 dataset was downloaded from the Gene Expression Omnibus (GEO) database, which contained 366 samples, including 200 heart failure samples and 166 non heart failure samples. The raw data was integrated to find differentially expressed genes (DEGs) and were further analyzed with bioinformatics analysis. Gene ontology (GO) and REACTOME enrichment analyses were performed via ToppGene; protein-protein interaction (PPI) networks of the DEGs was constructed based on data from the HiPPIE interactome database; modules analysis was performed; target gene - miRNA regulatory network and target gene - TF regulatory network were constructed and analyzed; hub genes were validated; molecular docking studies was performed. A total of 881 DEGs, including 442 up regulated genes and 439 down regulated genes were observed. Most of the DEGs were significantly enriched in biological adhesion, extracellular matrix, signaling receptor binding, secretion, intrinsic component of plasma membrane, signaling receptor activity, extracellular matrix organization and neutrophil degranulation. The top hub genes ESR1, PYHIN1, PPP2R2B, LCK, TP63, PCLAF, CFTR, TK1, ECT2 and FKBP5 were identified from the PPI network. Module analysis revealed that HF was associated with adaptive immune system and neutrophil degranulation. The target genes, miRNAs and TFs were identified from the target gene - miRNA regulatory network and target gene - TF regulatory network. Furthermore, receiver operating characteristic (ROC) curve analysis and RT-PCR analysis revealed that ESR1, PYHIN1, PPP2R2B, LCK, TP63, PCLAF, CFTR, TK1, ECT2 and FKBP5 might serve as prognostic, diagnostic biomarkers and therapeutic target for HF. The predicted targets of these active molecules were then confirmed. The current investigation identified a series of key genes and pathways that might be involved in the progression of HF, providing a new understanding of the underlying molecular mechanisms of HF.

## Introduction

Heart failure (HF) is a cardiovascular disease characterized by tachycardia, tachypnoea, pulmonary rales, pleural effusion, raised jugular venous pressure, peripheral oedema and hepatomegaly [1]. Morbidity and mortality linked with HF is a prevalent worldwide health problem holding a universal position as the leading cause of death [2]. The numbers of cases of HF are rising globally and it has become a key health issue. According to a survey, the prevalence HF is expected to exceed 50% of the global population [3]. Research suggests that modification in multiple genes and signaling pathways are associated in controlling the advancement of HF. However, a lack of investigation on the precise molecular mechanisms of HF development limits the treatment efficacy of the disease at present.

Previous study showed that HF was related to the expression of MECP2 [4] and RBM20 [5]. Toll-Like receptor signaling pathway [6] and activin type II receptor signaling pathway [7] were liable for progression of HF. More investigations are required to focus on treatments that enhance the outcome of patients with HF, to strictly make the diagnosis of the disease based on screening of biomarkers. These investigations can upgrade prognosis of patients by lowering the risk of advancement of HF and related complications. So it is essential to recognize the mechanism and find biomarkers with a good specificity and sensitivity.

The recent high-throughput RNA sequencing data has been widely employed to screen the differentially expressed genes (DEGs) between normal samples and HF samples in human beings, which makes it accessible for us to further explore the entire molecular alterations in HF at multiple levels involving DNA, RNA, proteins, epigenetic alterations, and metabolism [8]. However, there still exist obstacles to put these RNA seq data in application in clinic for the reason that the number of DEGs found by expression profiling by high throughput sequencing were massive and the statistical analyses were also too sophisticated [9–10]

In this study, first, we had chosen dataset GSE141910 from Gene Expression Omnibus (GEO) (http://www.ncbi.nlm.nih.gov/geo/) [11]. Second, we applied for limma tool in R software to obtain the differentially expressed genes (DEGs) in this dataset. Third, the ToppGene was used to analyze these DEGs including molecular function (MF), cellular component (CC), biological process (BP) and REACTOME pathways. Fourth, we established protein-protein interaction (PPI) network and then applied Cytotype PEWCC1 for module analysis of the DEGs which would identify some hub genes. Fifth, we established target gene - miRNA regulatory network and target gene - TF regulatory network. In addition, we further validated the hub genes by receiver operating characteristic (ROC) curve analysis and RT-PCR analysis. Finally, we performed molecular docking studies for over expressed hub genes. Results from the present investigation might provide new vision into potential prognostic and therapeutic targets for HF.

## Materials and Methods

### Data resource

Expression profiling by high throughput sequencing with series number GSE141910 based on platform GPL16791 was downloaded from the GEO database. The dataset of GSE141910 contained 200 heart failure samples and 166 non heart failure samples. It was downloaded from the GEO database in NCBI based on the platform of GPL16791 Illumina HiSeq 2500 (Homo sapiens).

### Identification of DEGs in HF

DEGs of dataset GSE141910 between HF groups and non heart failure groups were respectively analyzed using the limma package in R [12]. Fold changes (FCs) in the expression of individual genes were calculated and DEGs with P<0.05, |log FC| > 1.158 for up regulated genes and |log FC| < −0.83 for down regulated genes were considered to be significant. Hierarchical clustering and visualization were used by Heat-map package of R.

### Functional enrichment analysis

Gene Ontology (GO) analysis and REACTOME pathway analysis were performed to determine the functions of DEGs using the ToppGene (ToppFun) (https://toppgene.cchmc.org/enrichment.jsp) [13] GO terms (http://geneontology.org/) [14] included biological processes (BP), cellular components (CC) and molecular functions (MF) of genomic products.

REACTOME (https://reactome.org/) [15] analyzes pathways of important gene products. ToppGene is a bioinformatics database for analyzing the functional interpretation of lists of proteins and genes. The cutoff value was set to P<0.05.

### Protein–protein interaction network construction and module screening

PPI networks are used to establish all protein coding genes into a massive biological network that serves an advance compassionate of the functional system of the proteome [16]. The HiPPIE interactome (https://cbdm.uni-mainz.de/hippie/) [17] database furnish information regarding predicted and experimental interactions of proteins. In the current investigation, the DEGs were mapped into the HiPPIE interactome database to find significant protein pairs with a combined score of >0.4. The PPI network was subsequently constructed using Cytoscape software, version 3.8.2 (www.cytoscape.org) [18]. The nodes with a higher node degree [19], higher betweenness centrality [20], higher stress centrality [21] and higher closeness centrality [22] were considered as hub genes. Additionally, cluster analysis for identifying significant function modules with a degree cutoff >2 in the PPI network was performed using the PEWCC1 (http://apps.cytoscape.org/apps/PEWCC1) [23] in Cytoscape.

### Target gene - miRNA regulatory network construction

The miRNet database (https://www.mirnet.ca/) [24] contains information on miRNA and the regulated genes. Using information collected from the miRNet database, hub genes were matched with their associated miRNA. The target gene - miRNA regulatory network then was constructed using Cytoscape software.

### Target gene - TF regulatory network construction

The NetworkAnalyst database (https://www.networkanalyst.ca/) [25] contains information on TF and the regulated genes. Using information collected from the NetworkAnalyst database, hub genes were matched with their associated TF. The target gene - TF regulatory network then was constructed using Cytoscape software.

### Receiver operating characteristic (ROC) curve analysis

Then ROC curve analysis was implement to classify the sensitivity and specificity of the hub genes for HF diagnosis and we investigated how large the area under the curve (AUC) was by using the statistical package pROC in R software [26].

### RT-PCR analysis

Total RNA was isolated from H9c2 (ATCC® CRL-1446) and VH2 (ATCC® CCL-140) using the TRI Reagent® (Sigma, USA). cDNA was synthesized using 2.0 μ of total RNA with the Reverse transcription cDNA kit (Thermo Fisher Scientific, Waltham, MA, USA). The 7 Flex real-time PCR system (Thermo Fisher Scientific, Waltham, MA, USA) was employed to detect the relative mRNA expression. The relative expression levels were determined by the 2-ΔΔCt method and normalized to internal control beta-actin [27]. All RT-PCR reactions were performed in triplicate. The primers used to explore mRNA expression of ten hub genes were listed in Table 1.

**Table 1.**
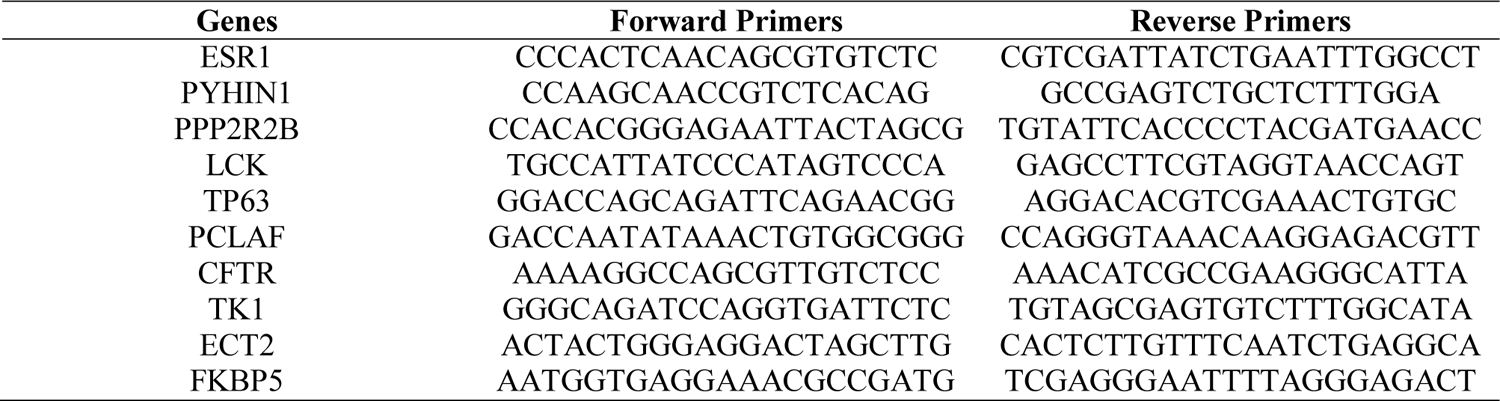
The sequences of primers for quantitative RT-PCR

### Identification of Candidate Small Molecules

SYBYL-X 2.0 perpetual drug design software has been used for surflex-docking studies of the designed novel molecules and the standard on over expressed genes of PDB protein. Using ChemDraw Software, all designed molecules and standards were sketched, imported and saved using open babel free software in sdf. template. The protein of over expressed genes of ESR1, LCK, PPP2R2B, TP63 and their co-crystallised protein of PDB code 4PXM, 1KSW, 2HV7, 3VD8 and 6RU6 were extracted from Protein Data Bank [28–31]. Optimizations of the designed molecules were performed by standard process by applying Gasteiger Huckel (GH) charges together with the TRIPOS force field. In addition, energy minimization was achieved using MMFF94s and MMFF94 algorithm methods. The preparation of the protein was done after protein incorporation. The co-crystallized ligand and all water molecules have been eliminated from the crystal structure; more hydrogen’s were added and the side chain was set, TRIPOS force field was used for the minimization of structure. The interaction efficiency of the compounds with the receptor was expressed in kcal / mol units by the Surflex-Dock score. The best location was integrated into the molecular region by the interaction between the protein and the ligand. Using Discovery Studio Visualizer, the visualisation of ligand interaction with receptor is performed.

## Results

### Identification of DEGs in HF

We identified 881 DEGs in the GSE141910 dataset using the limma package in R. Based on the limma analysis, using the adj P val <0.05, |log FC| > 1.158 for up regulated genes and |log FC| < −0.83 for down regulated genes criteria, a total of 881 DEGs were identified, consisting of 442 genes were up regulated and 439 genes were down regulated. The DEGs are listed in Table 2. The volcano plot for DEGs is illustrated in Fig. 1. Fig. 2 is the hierarchical clustering heat-map.

**Fig. 1.**
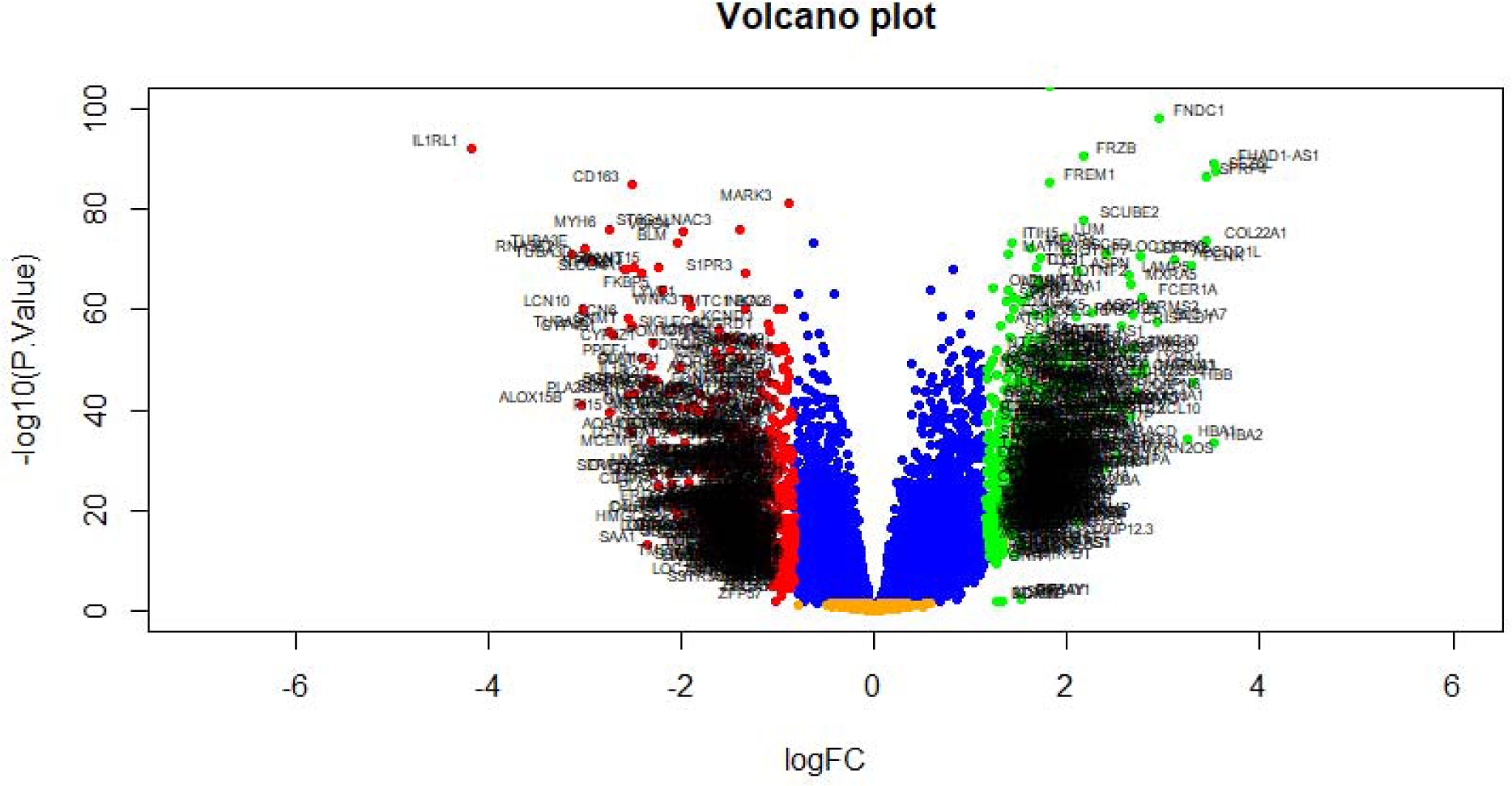
Volcano plot of differentially expressed genes. Genes with a significant change of more than two-fold were selected. Green dot represented up regulated significant genes and red dot represented down regulated significant genes.

**Fig. 2.**
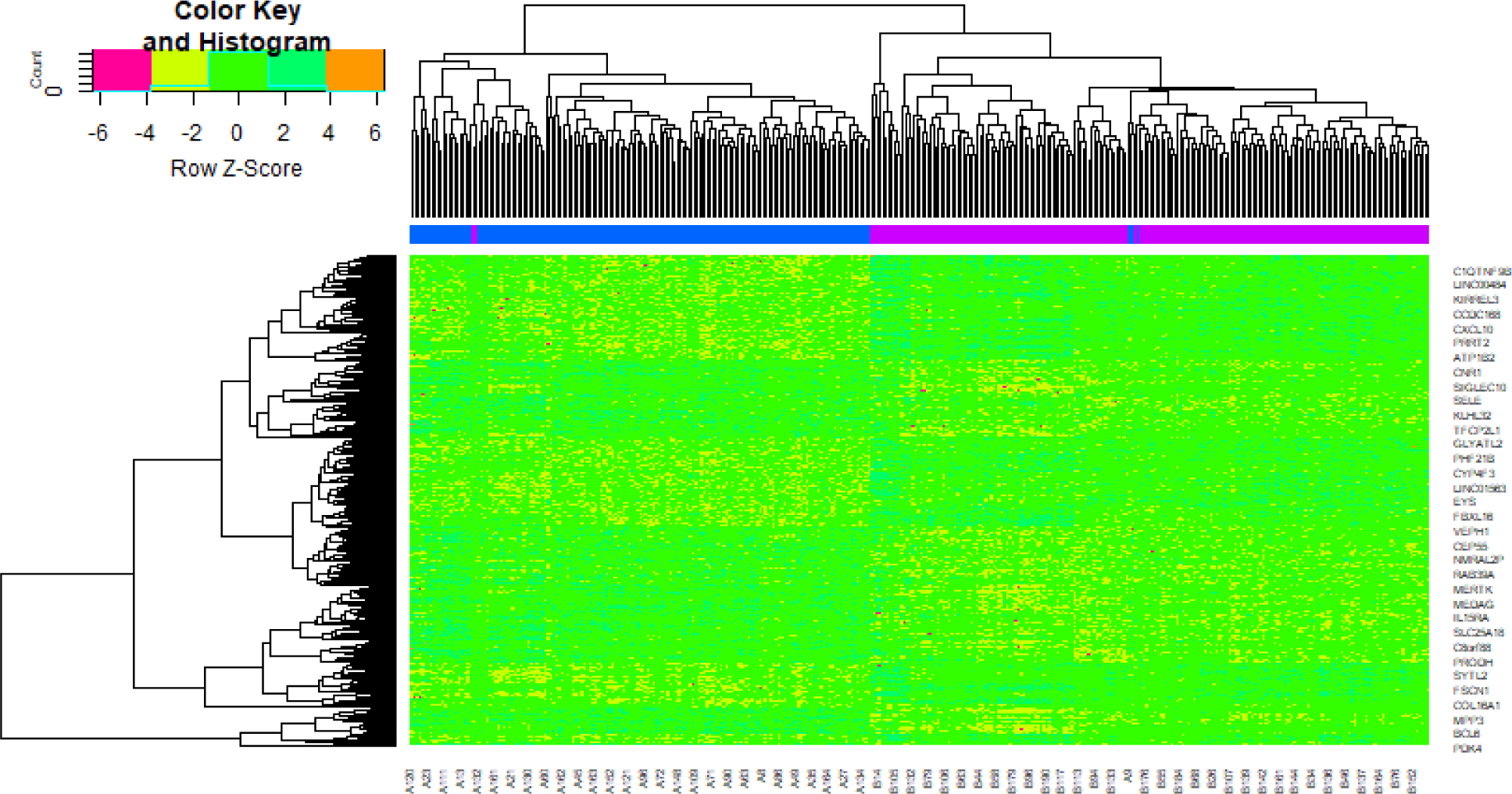
Heat map of differentially expressed genes. Legend on the top left indicate log fold change of genes. (A1 – A200= heart failure samples; B1 – B166 = non heart failure samples)

**Table 2.**
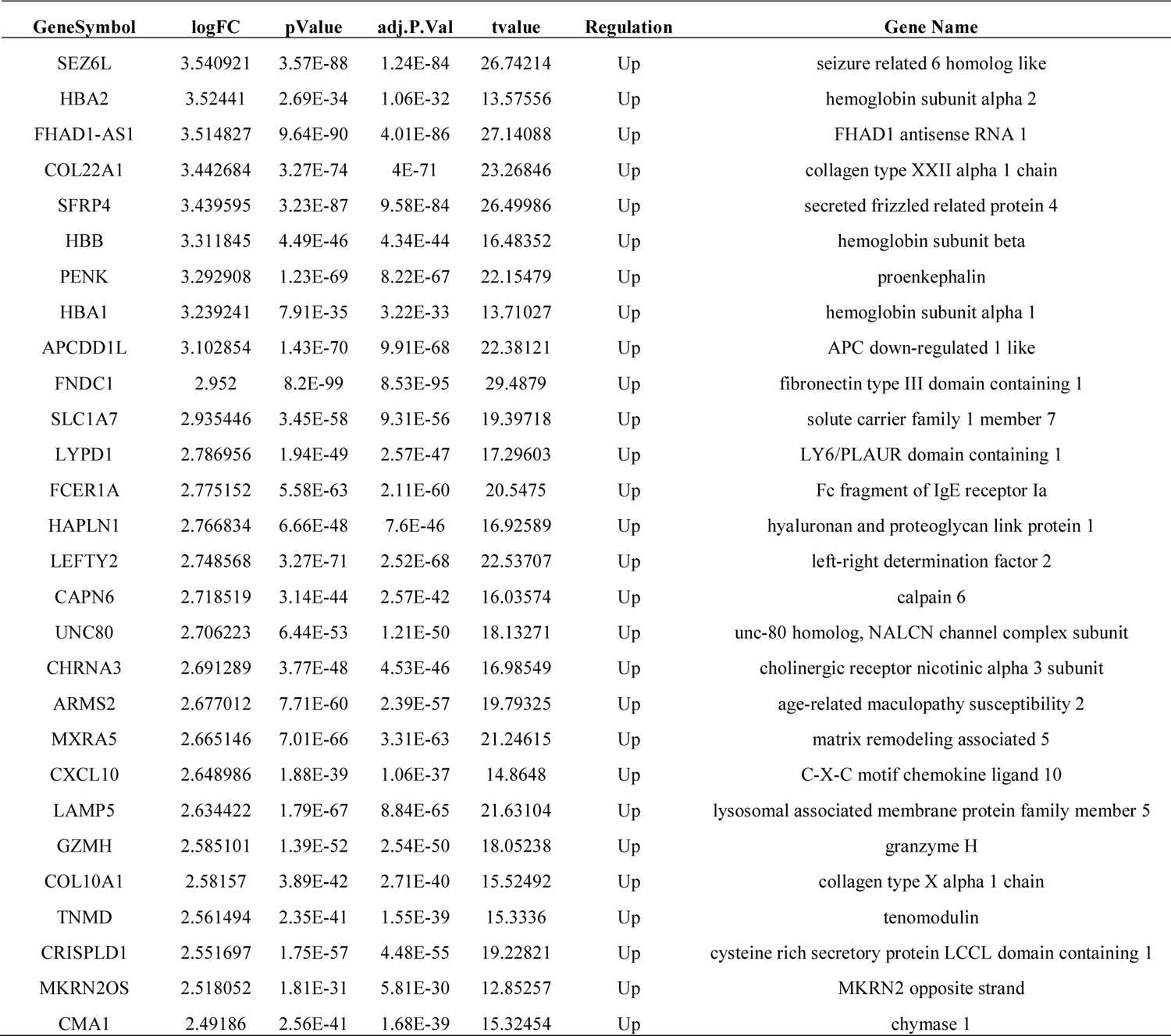
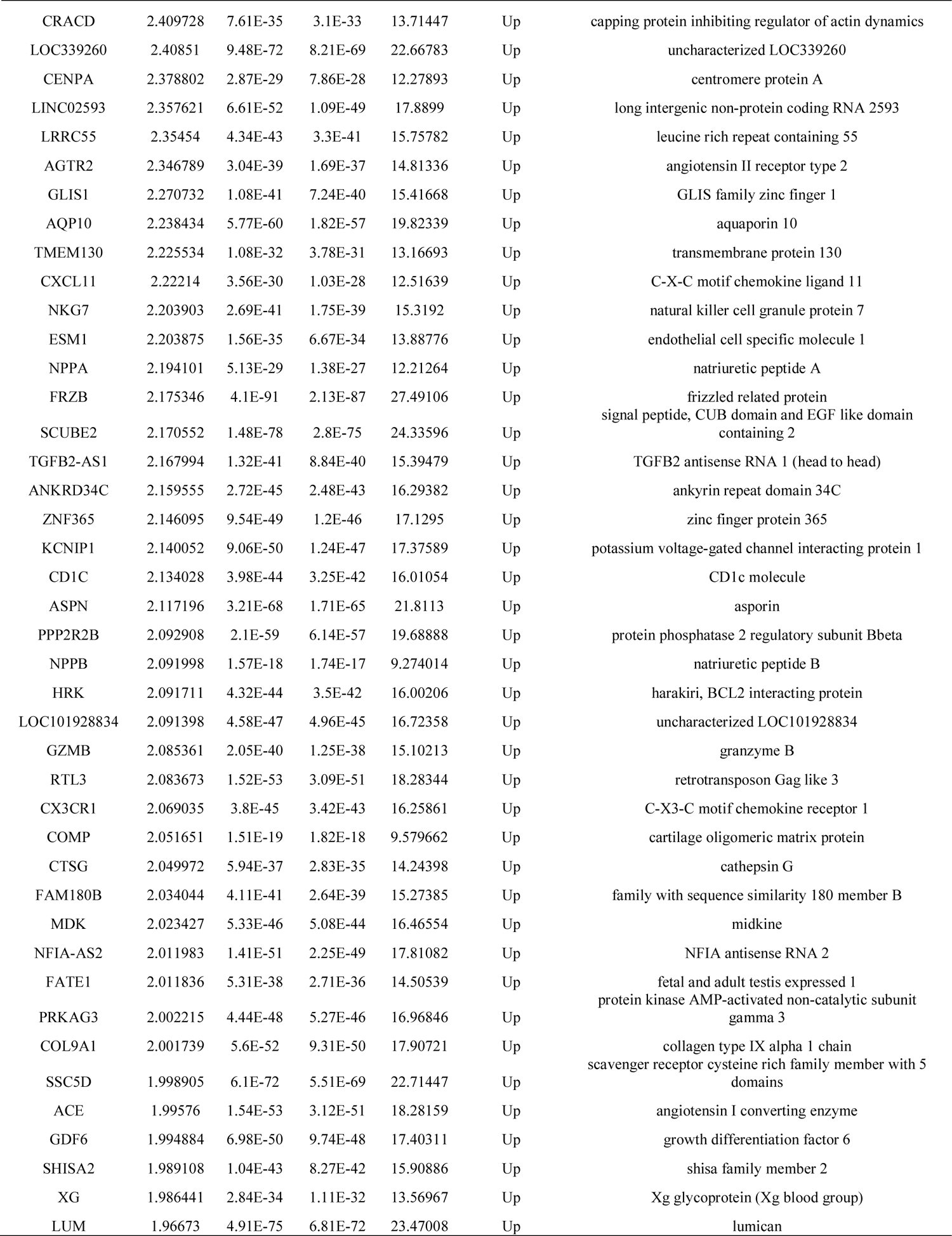
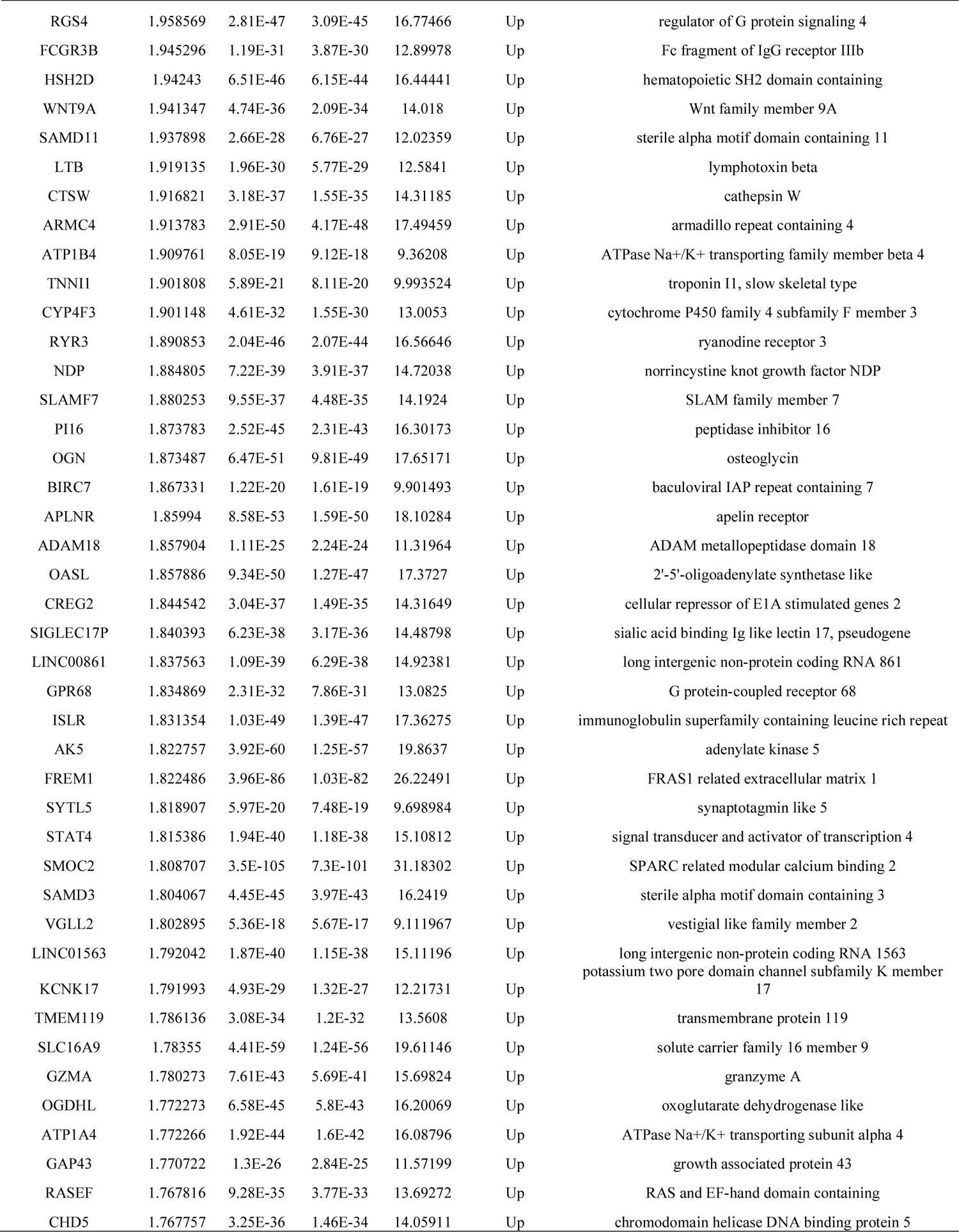
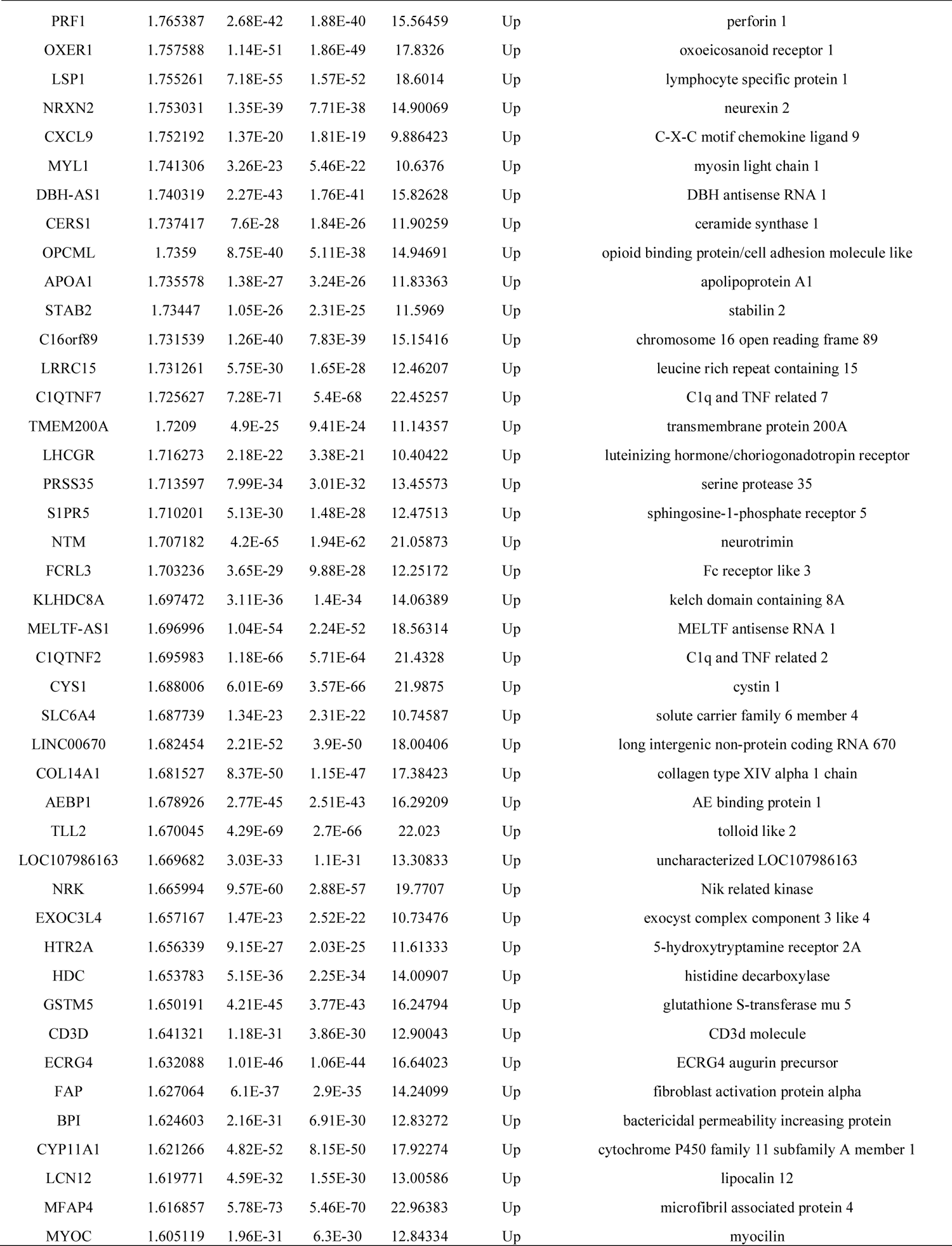
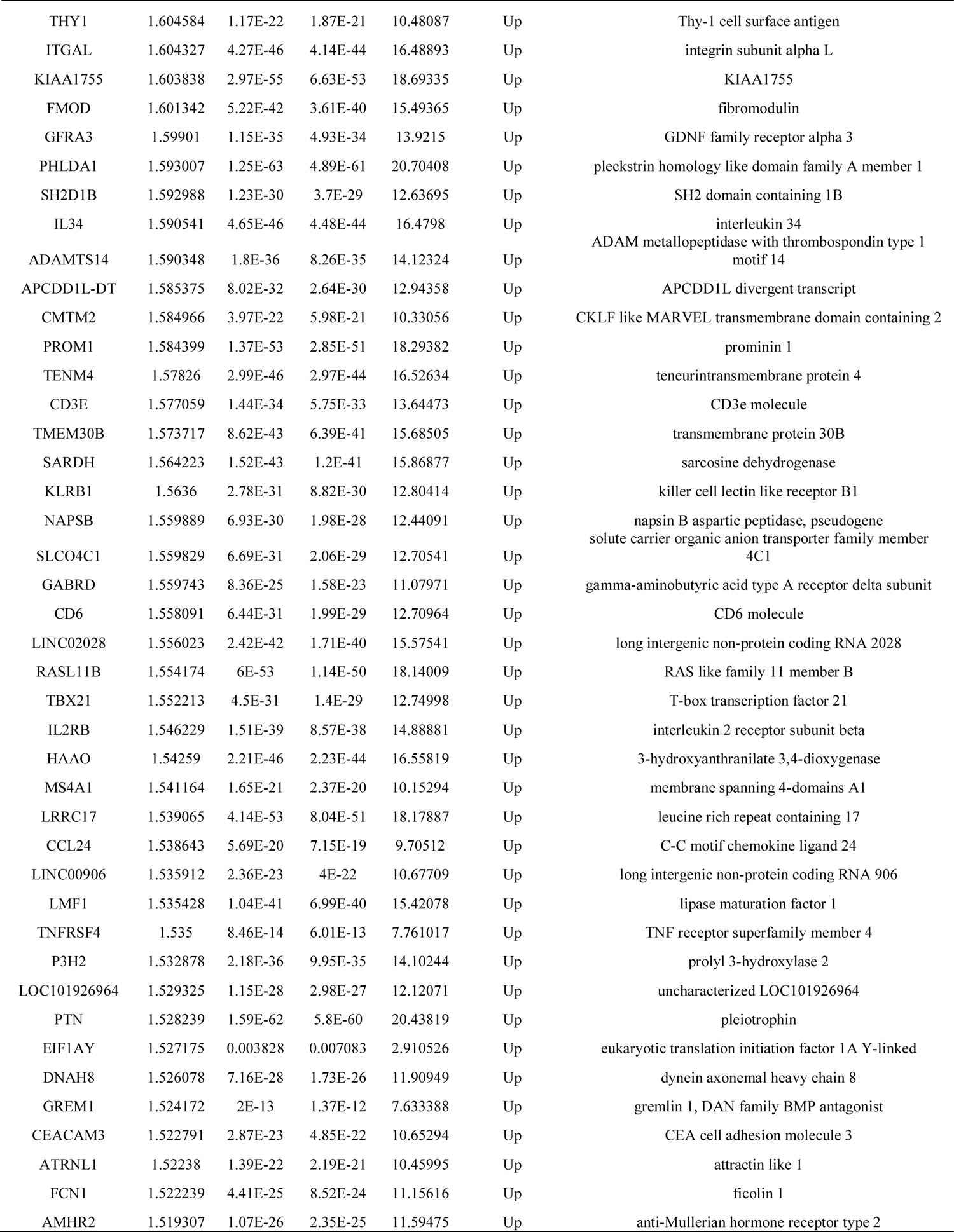
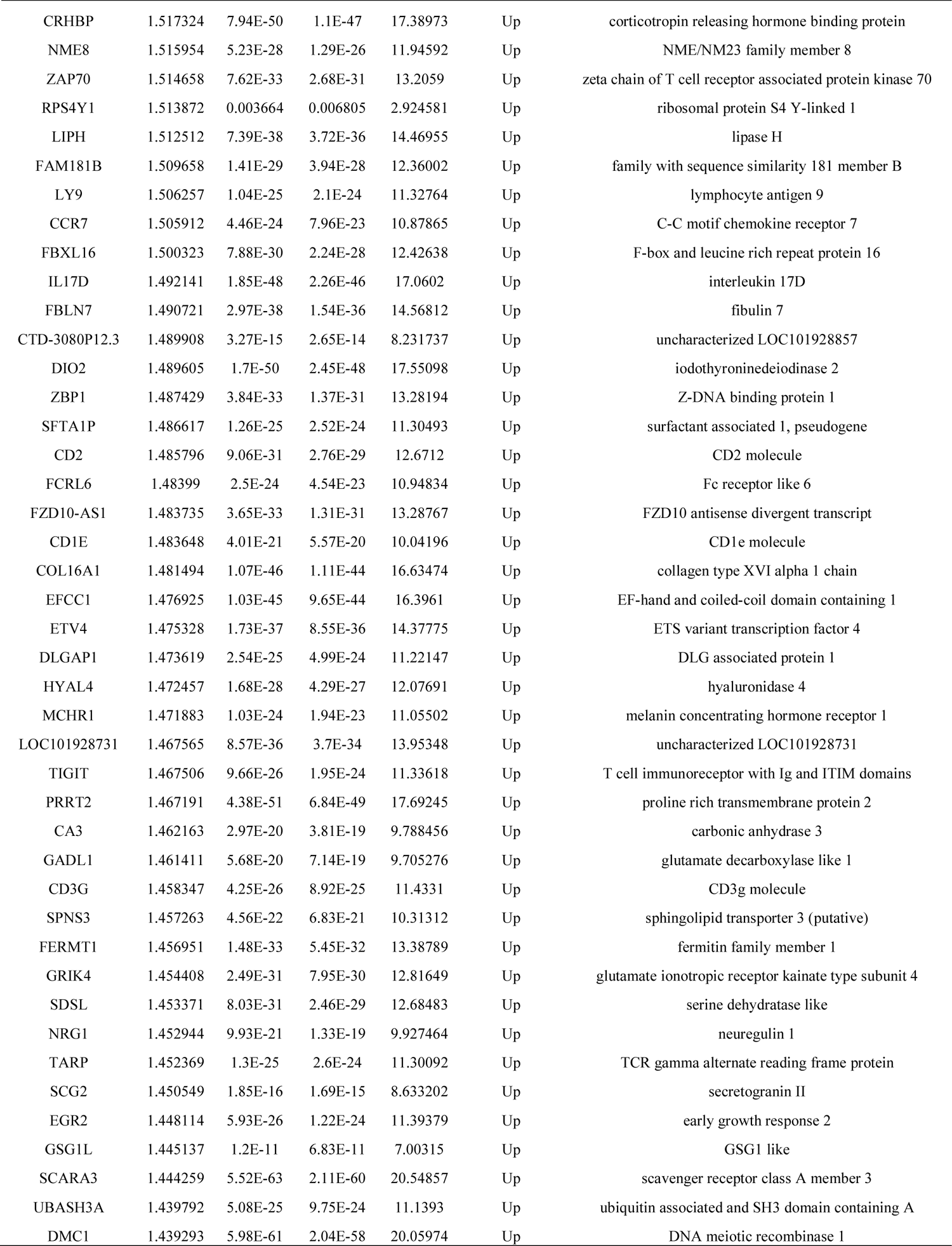
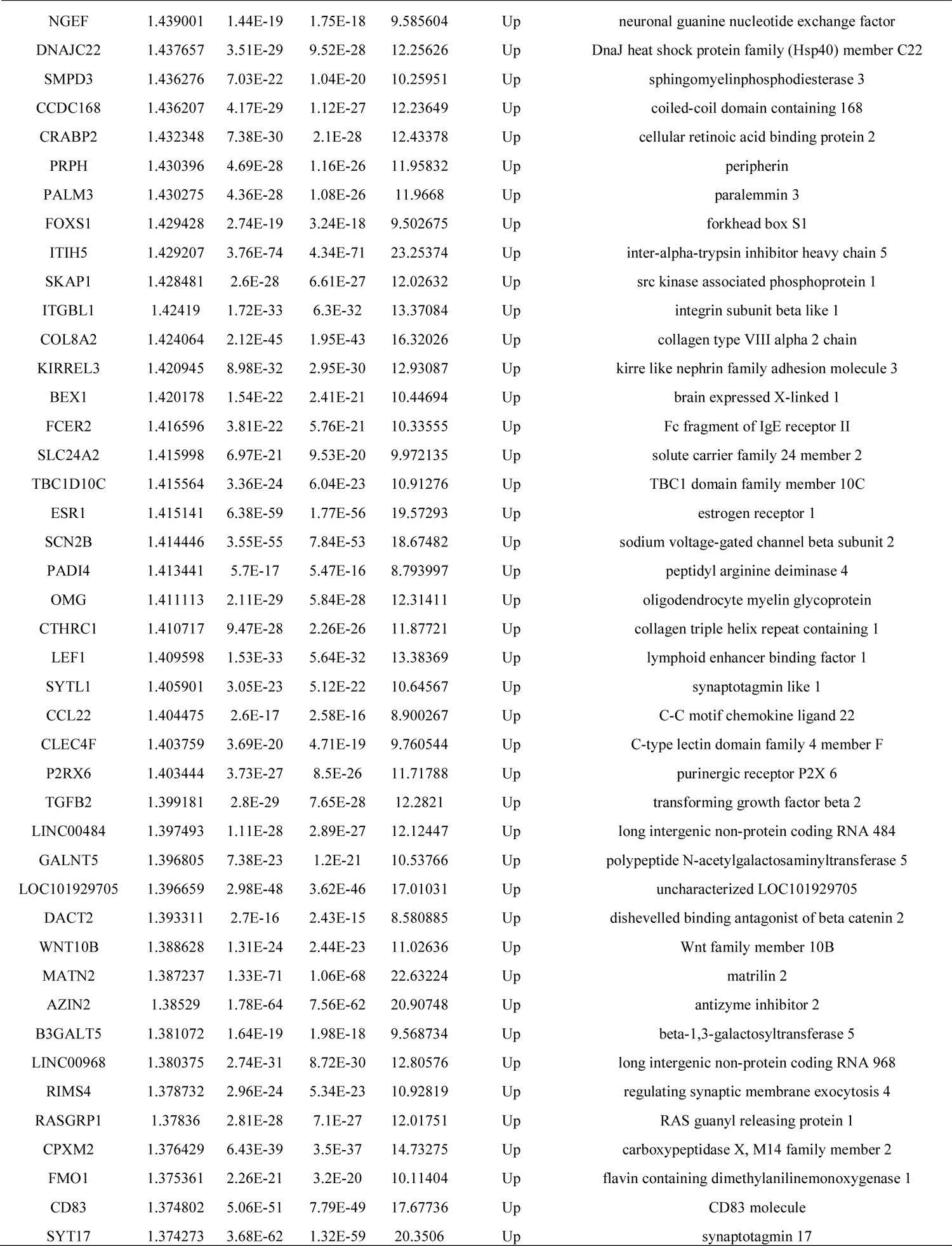
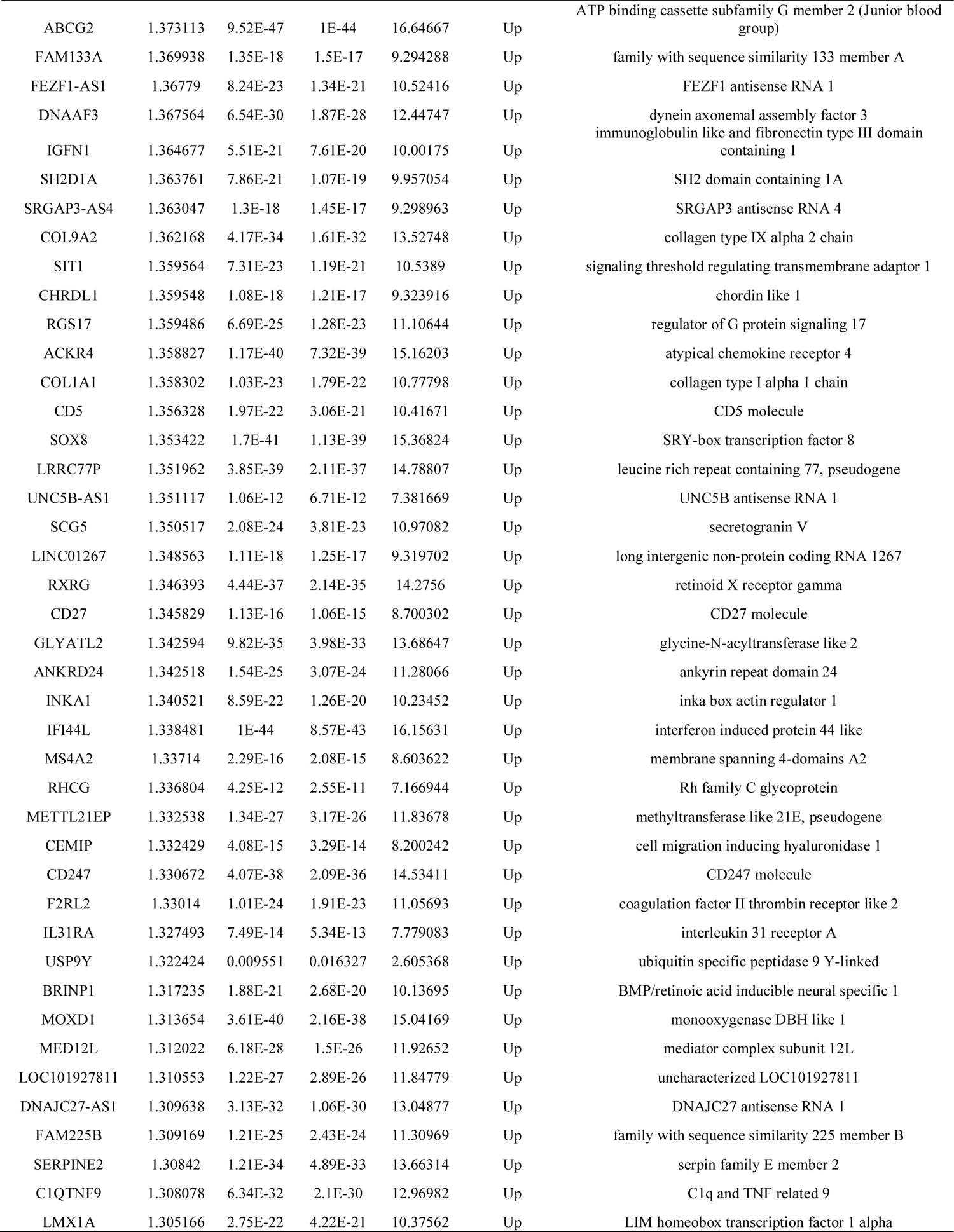
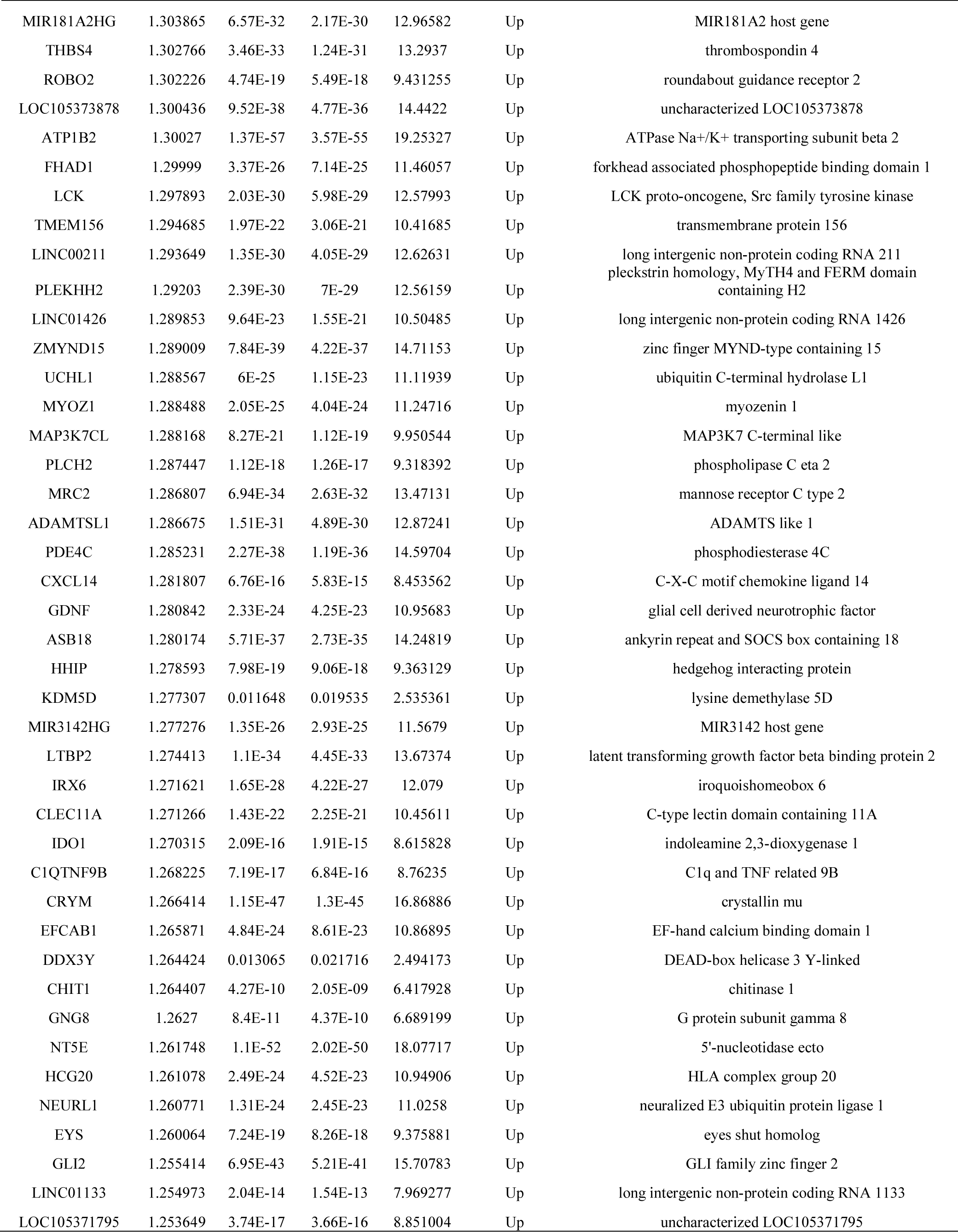
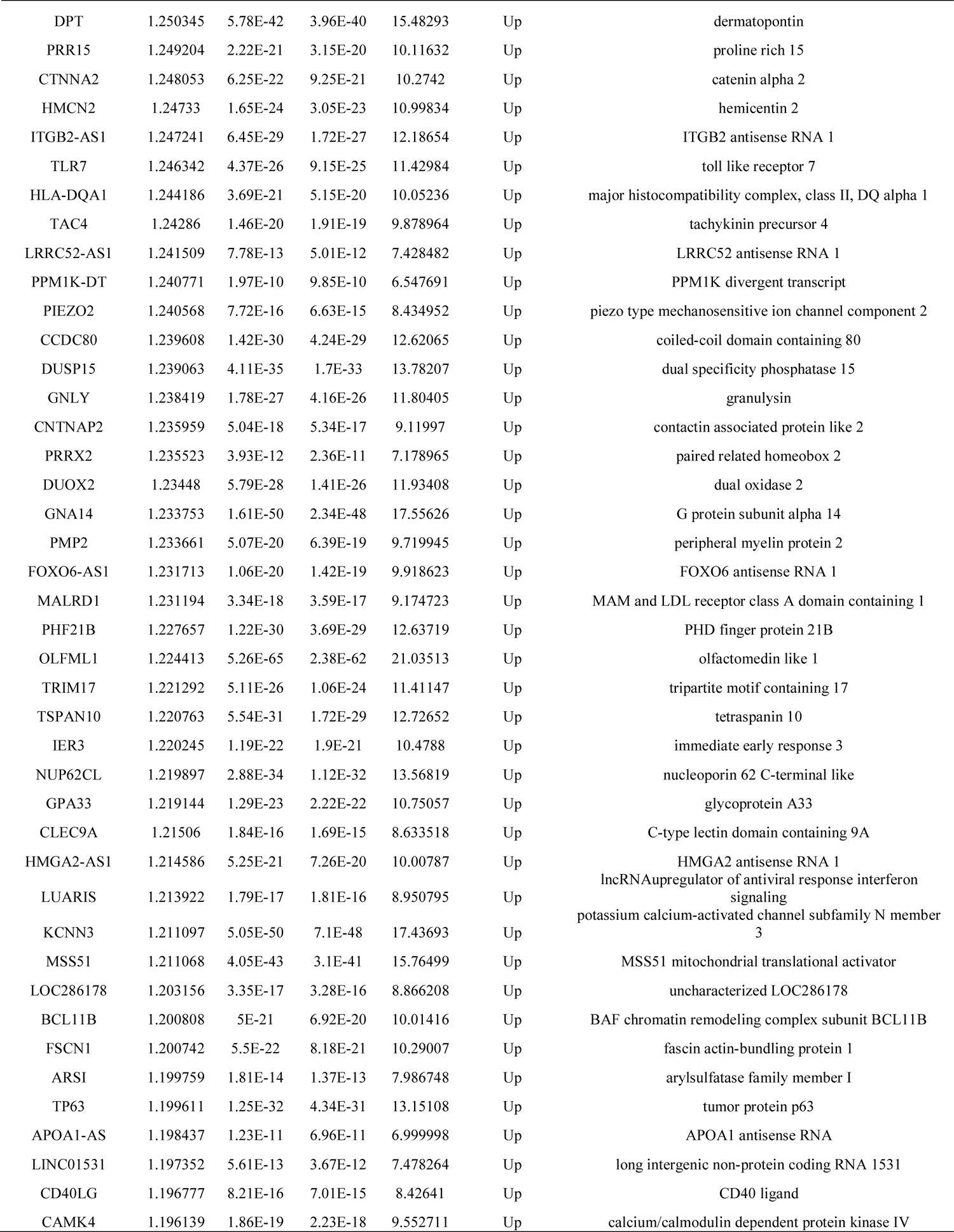
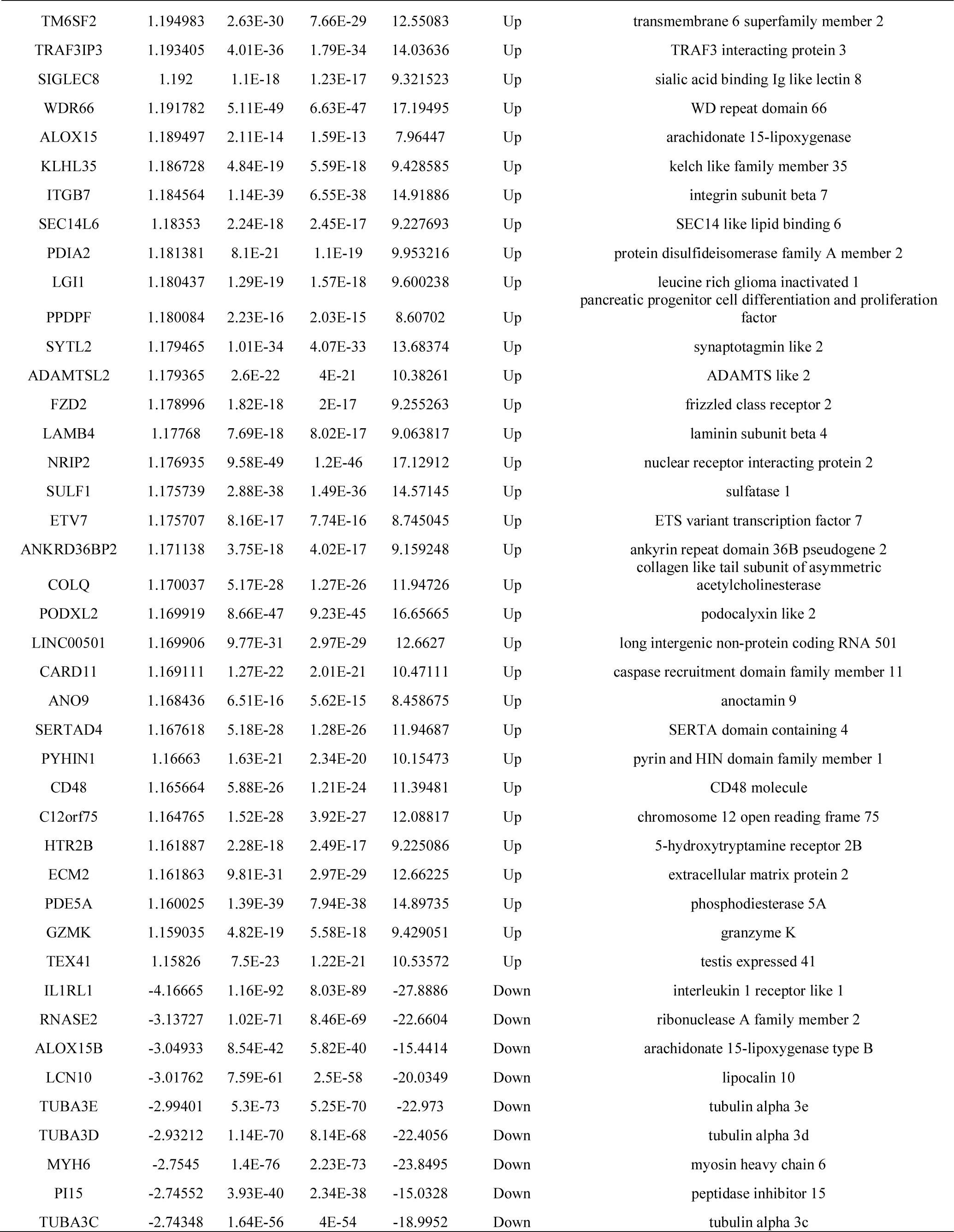
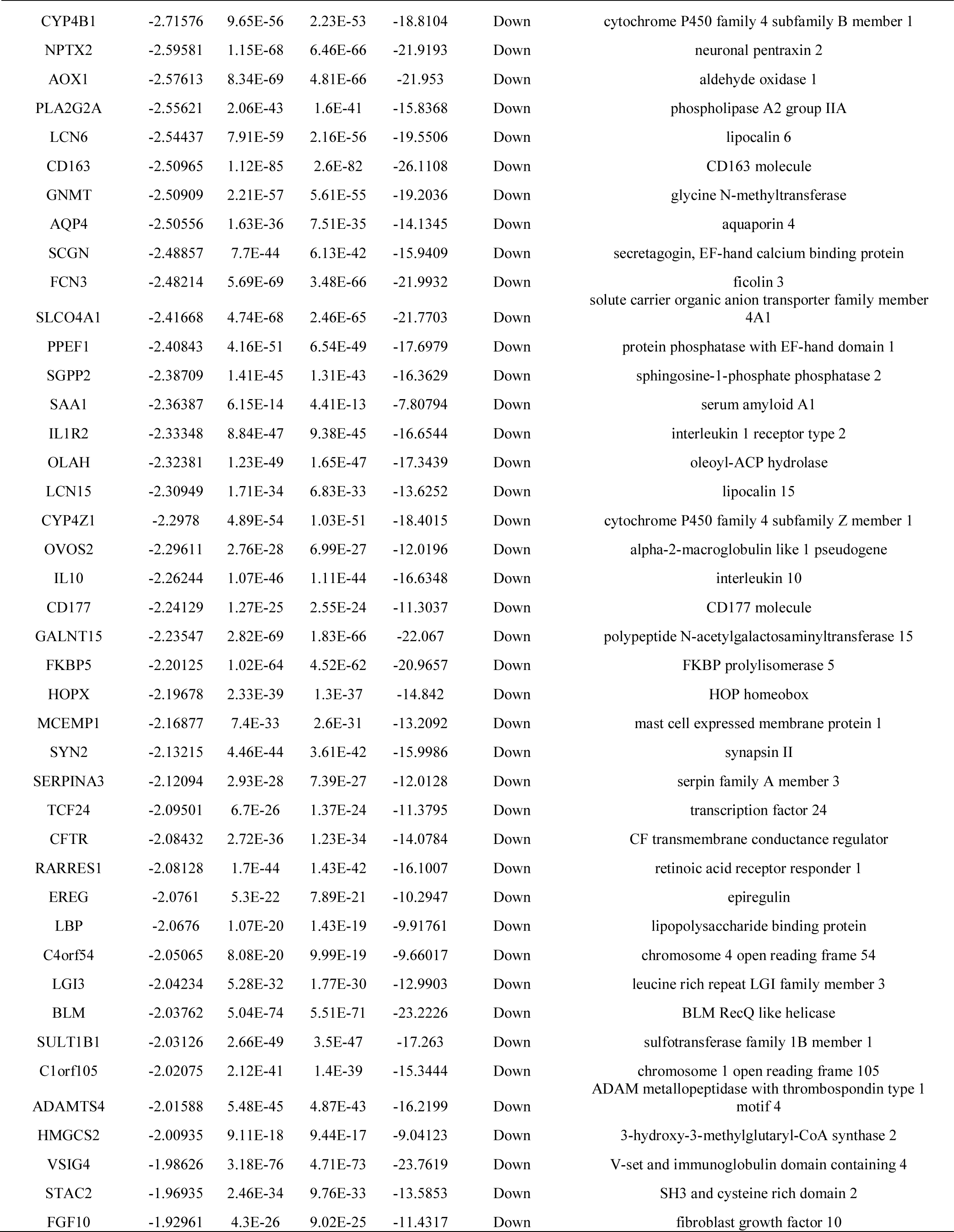
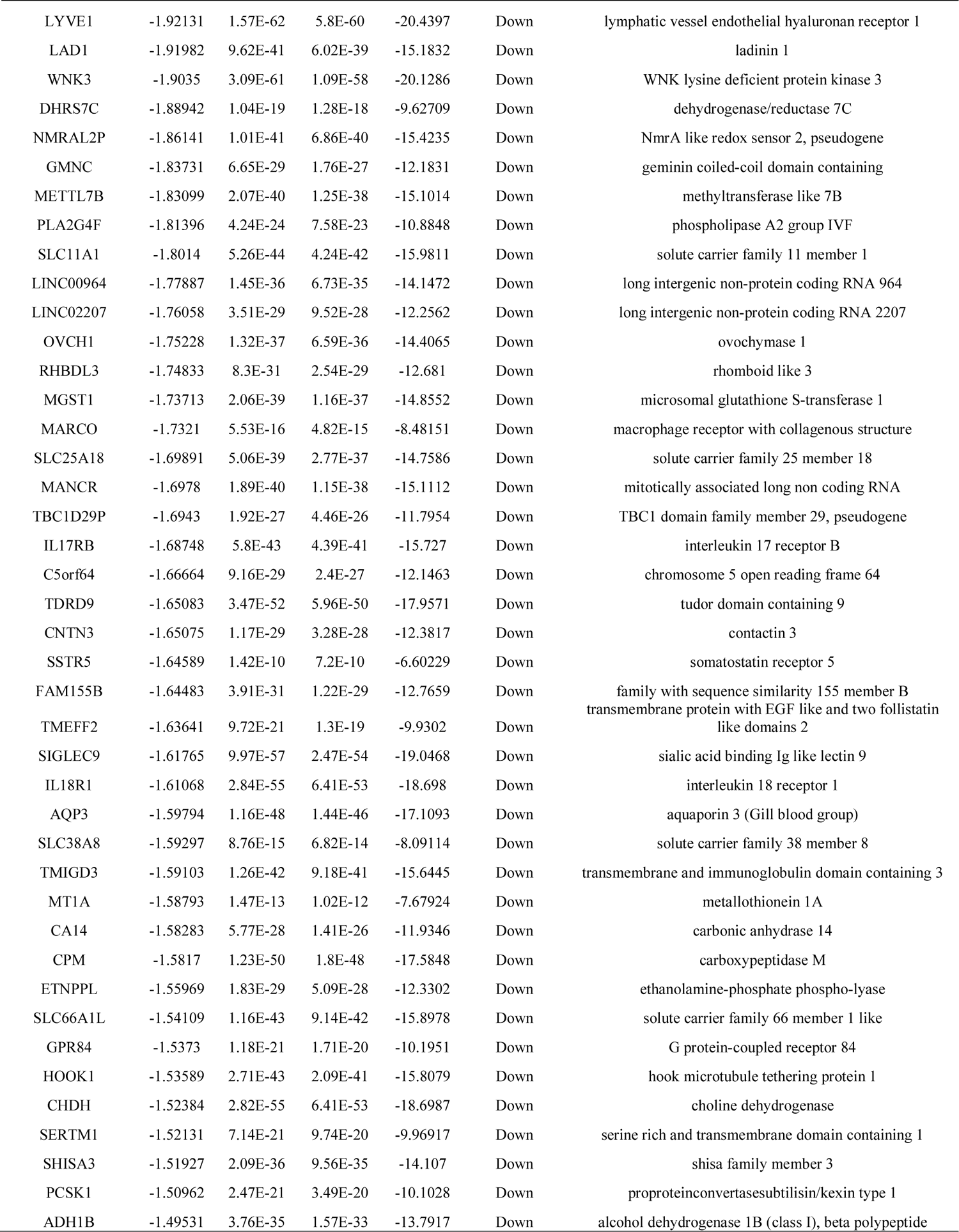
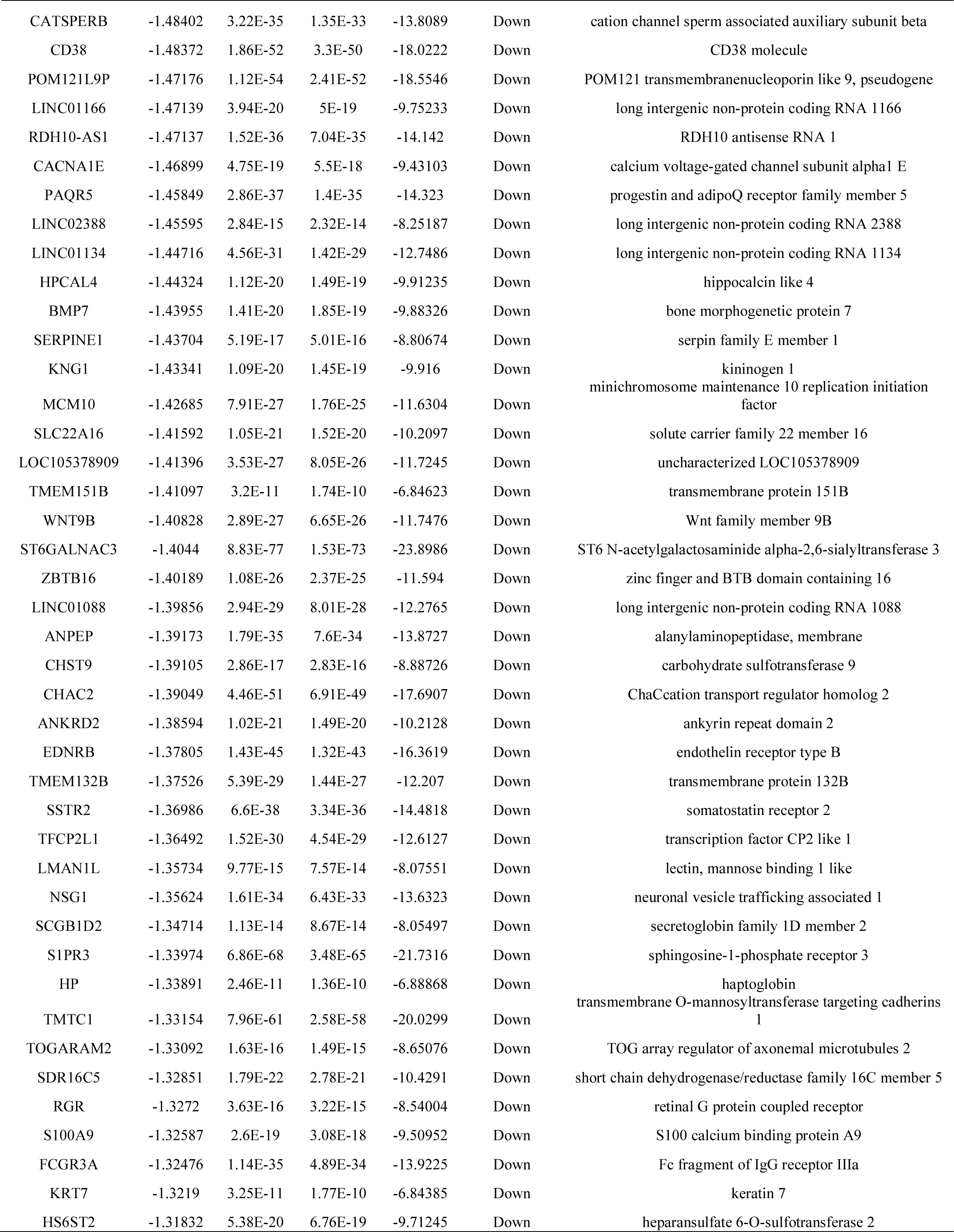
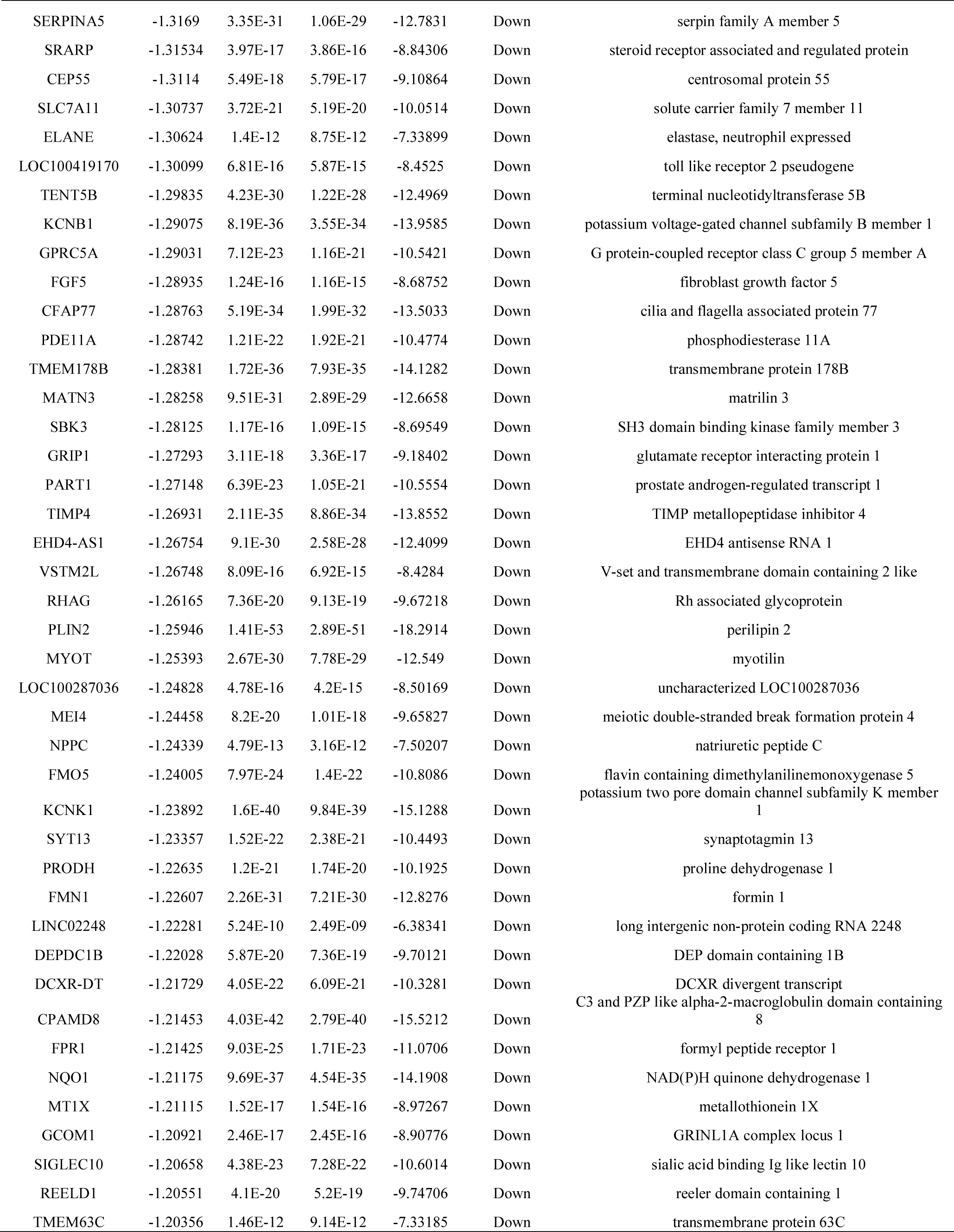
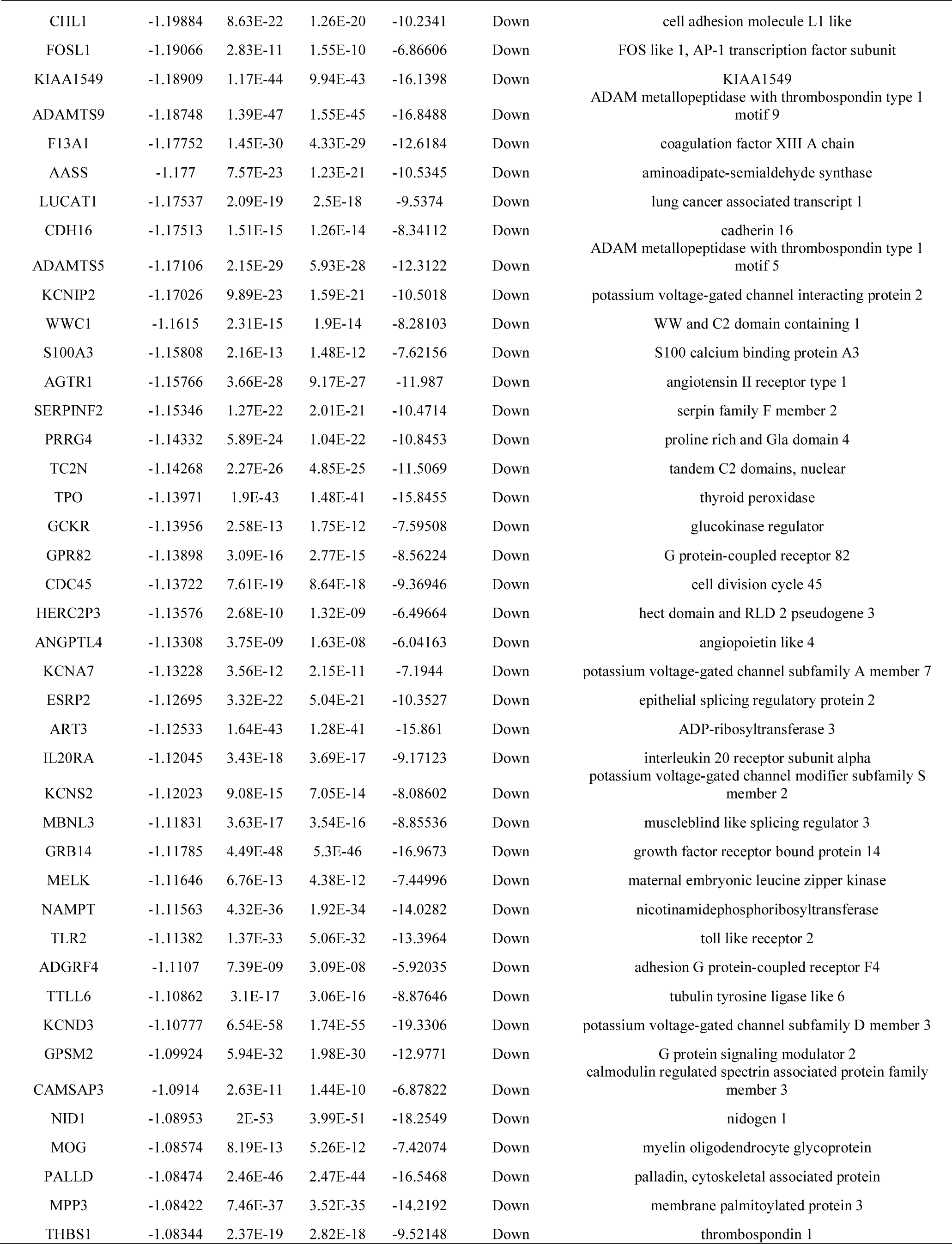
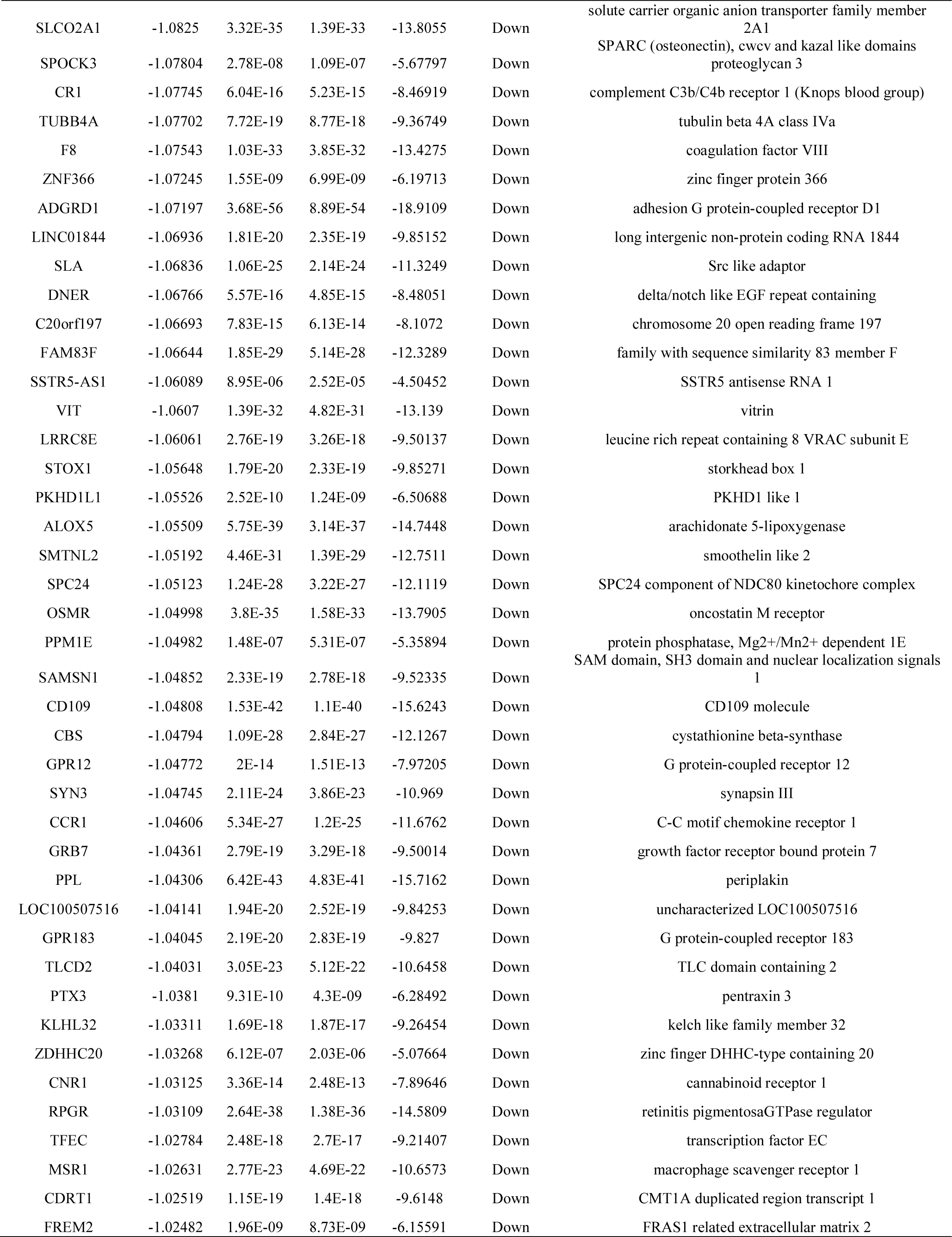
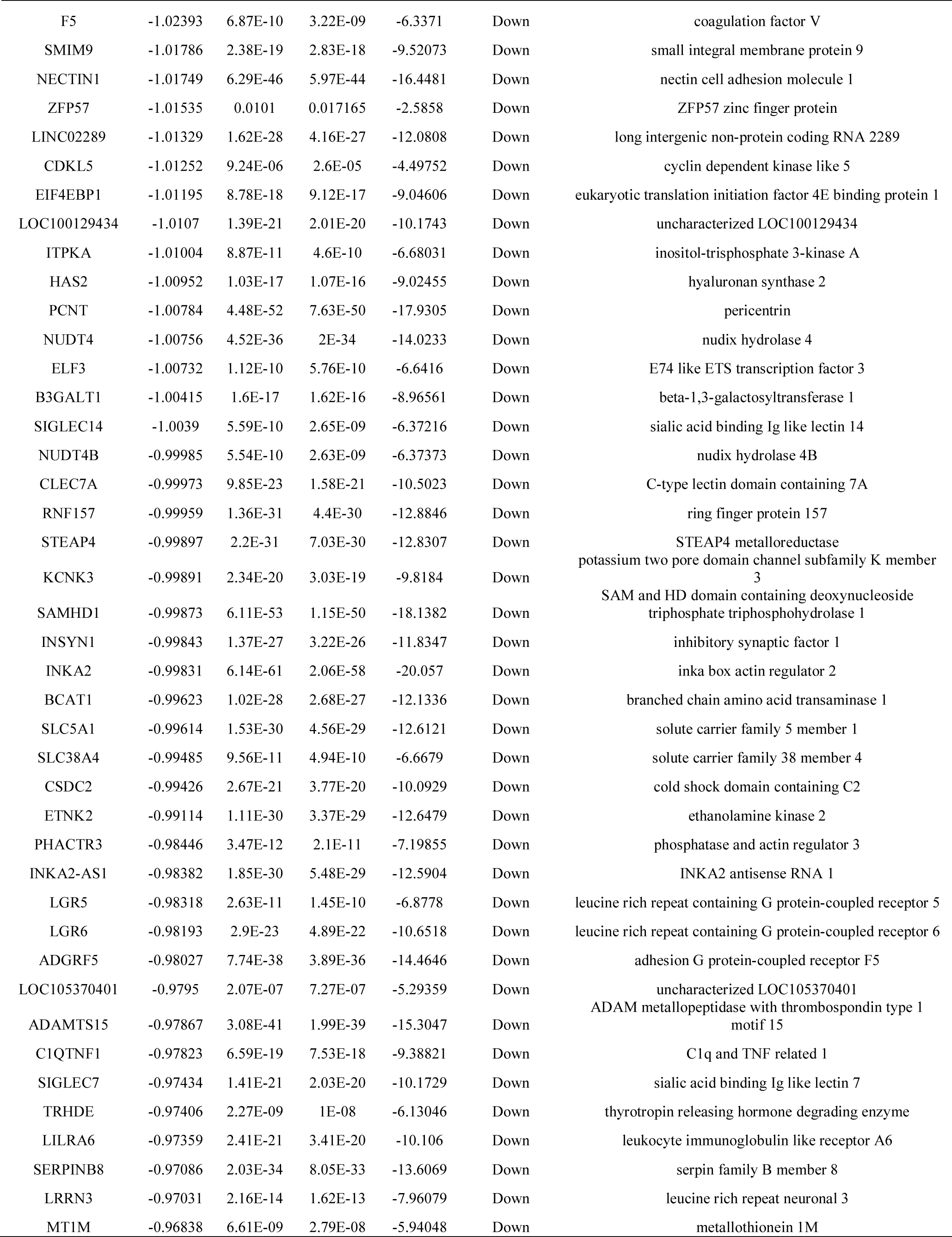
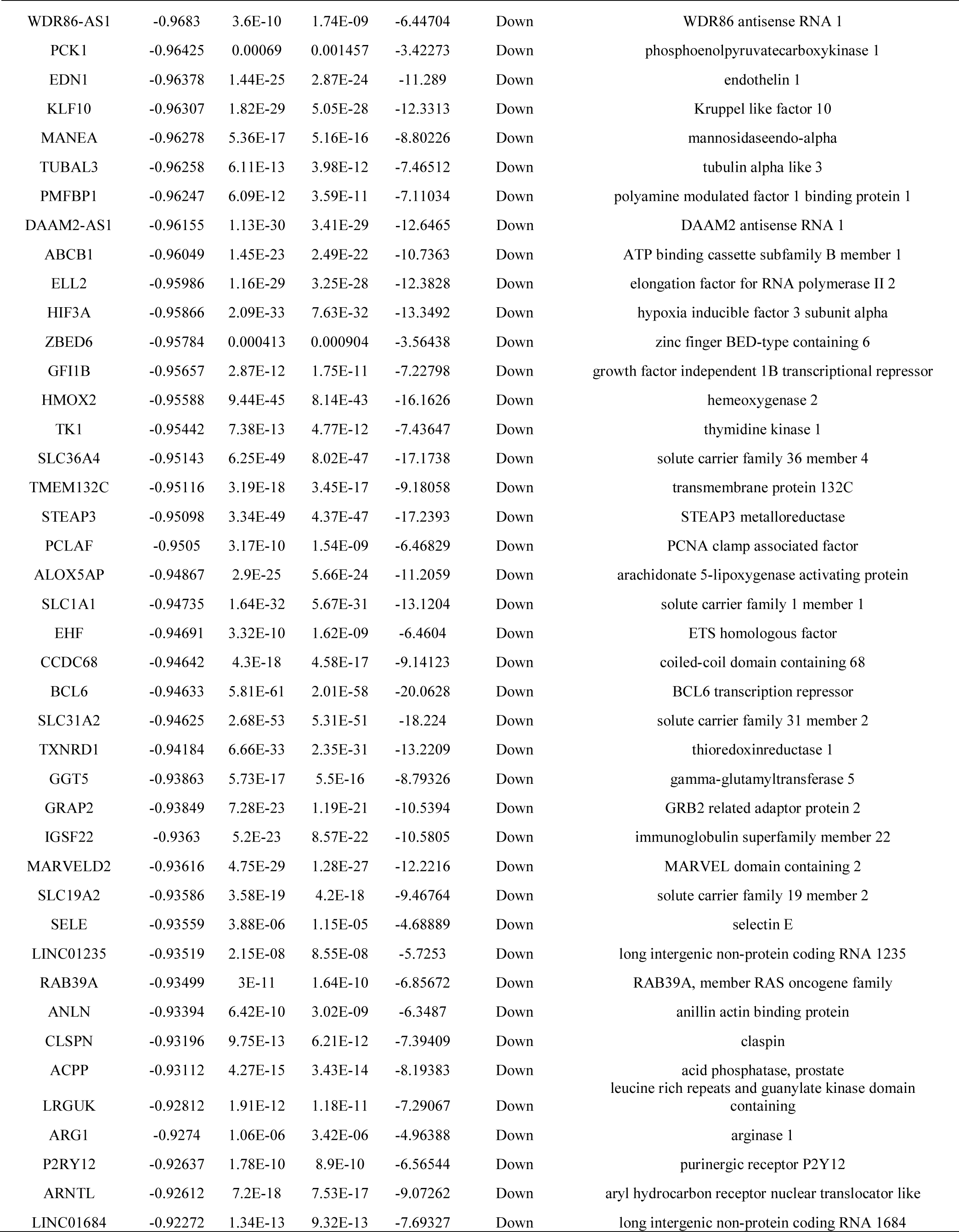
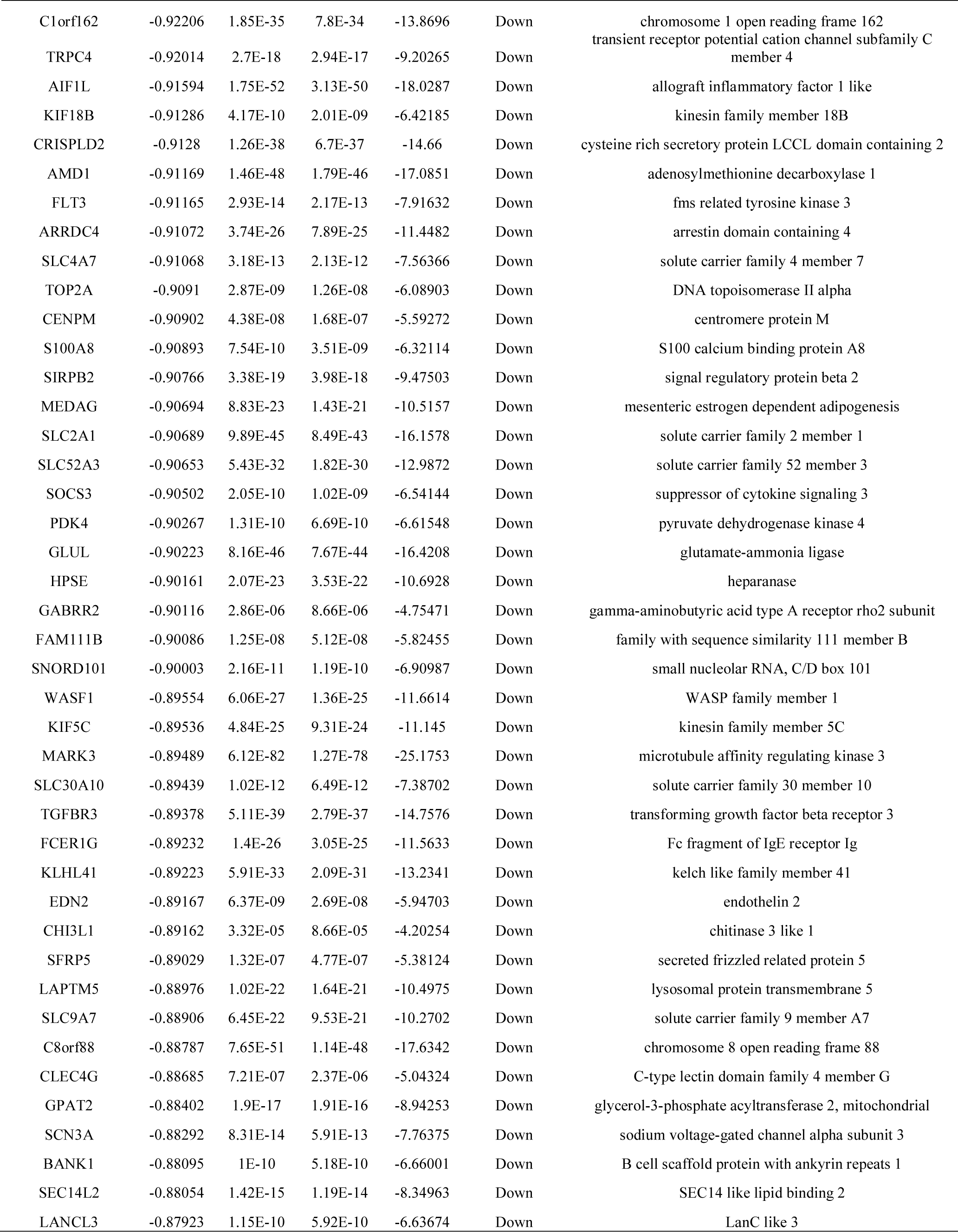
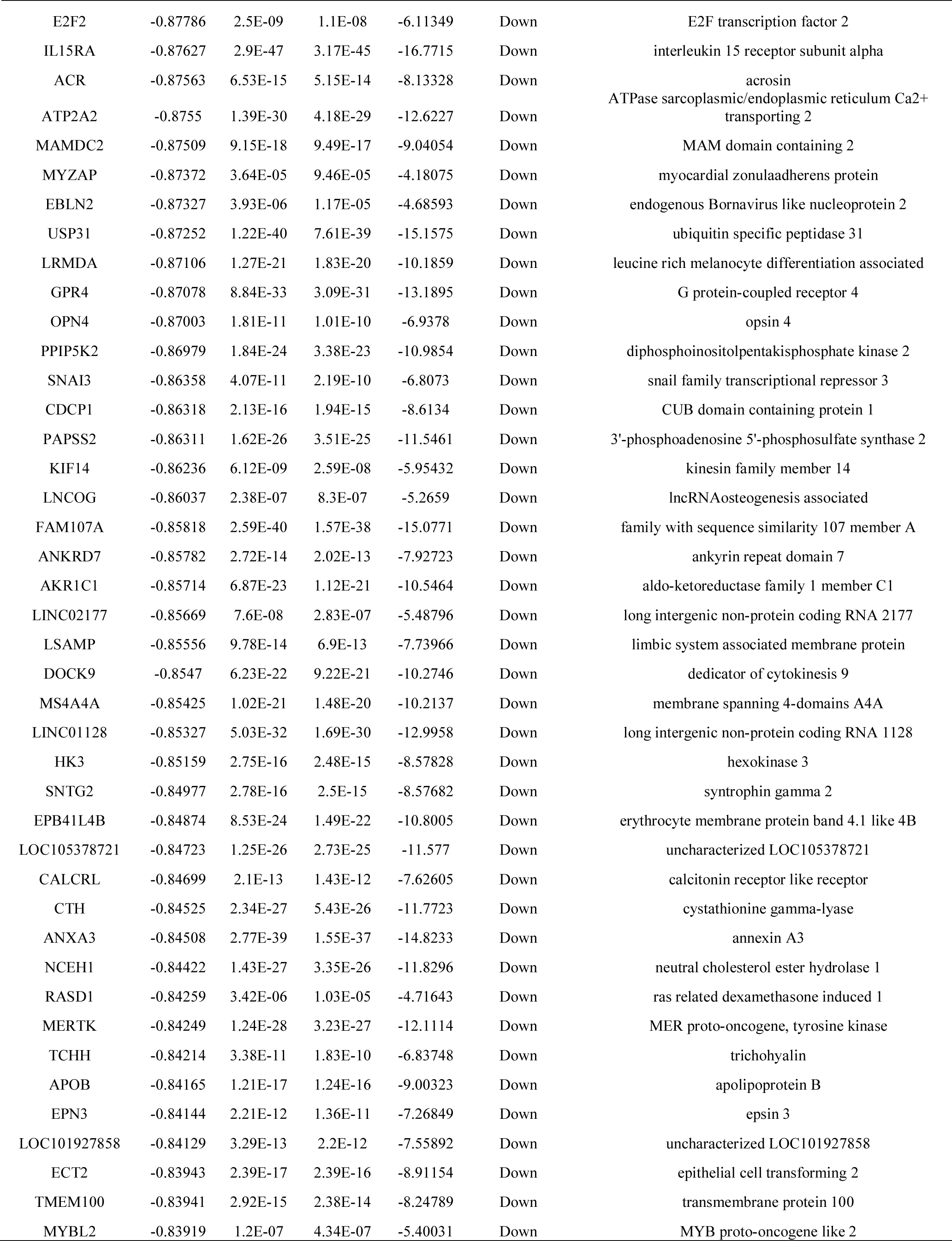
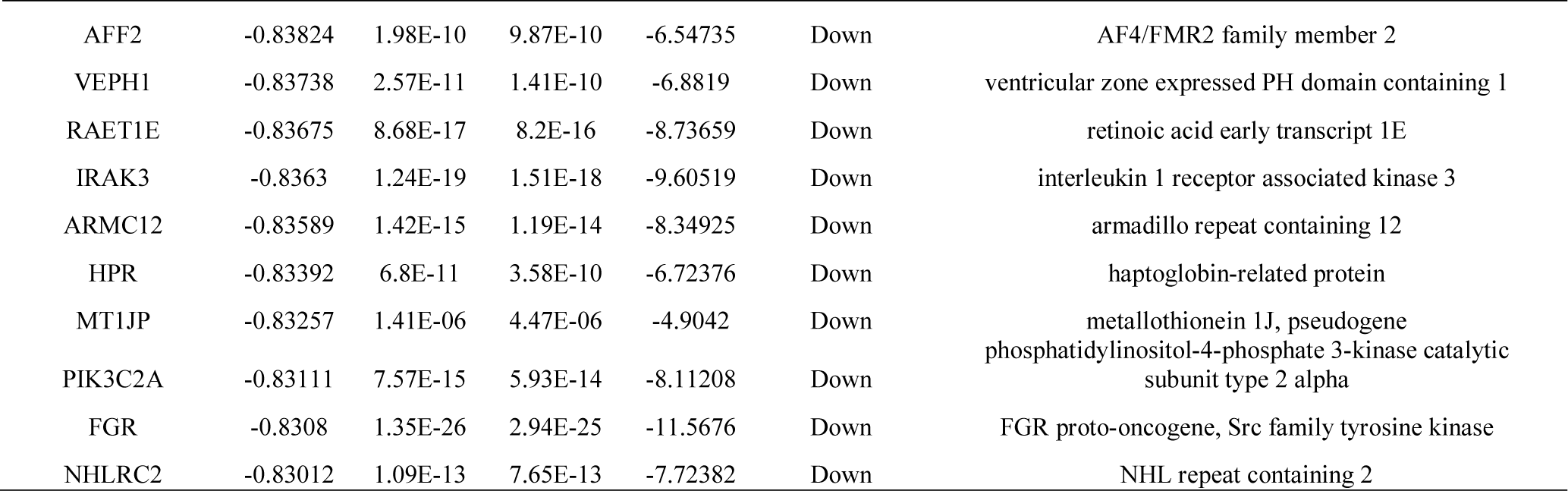
The statistical metrics for key differentially expressed genes (DEGs)

### Functional enrichment analysis

Results of GO analysis showed that the up regulated genes were significantly enriched in BP, CC, and MF, including biological adhesion, regulation of immune system process, extracellular matrix, cell surface, signaling receptor binding and molecular function regulator (Table 3); the down regulated genes were significantly enriched in BP, CC, and MF, including secretion, defense response, intrinsic component of plasma membrane, whole membrane, signaling receptor activity and molecular transducer activity (Table 3). Pathway analysis showed that the up regulated genes were significantly enriched in extracellular matrix organization and immunoregulatory interactions between a lymphoid and a non-lymphoid cell (Table 4); the down regulated genes were significantly enriched in neutrophil degranulation and SLC-mediated transmembrane transport (Table 4).

**Table 3.**
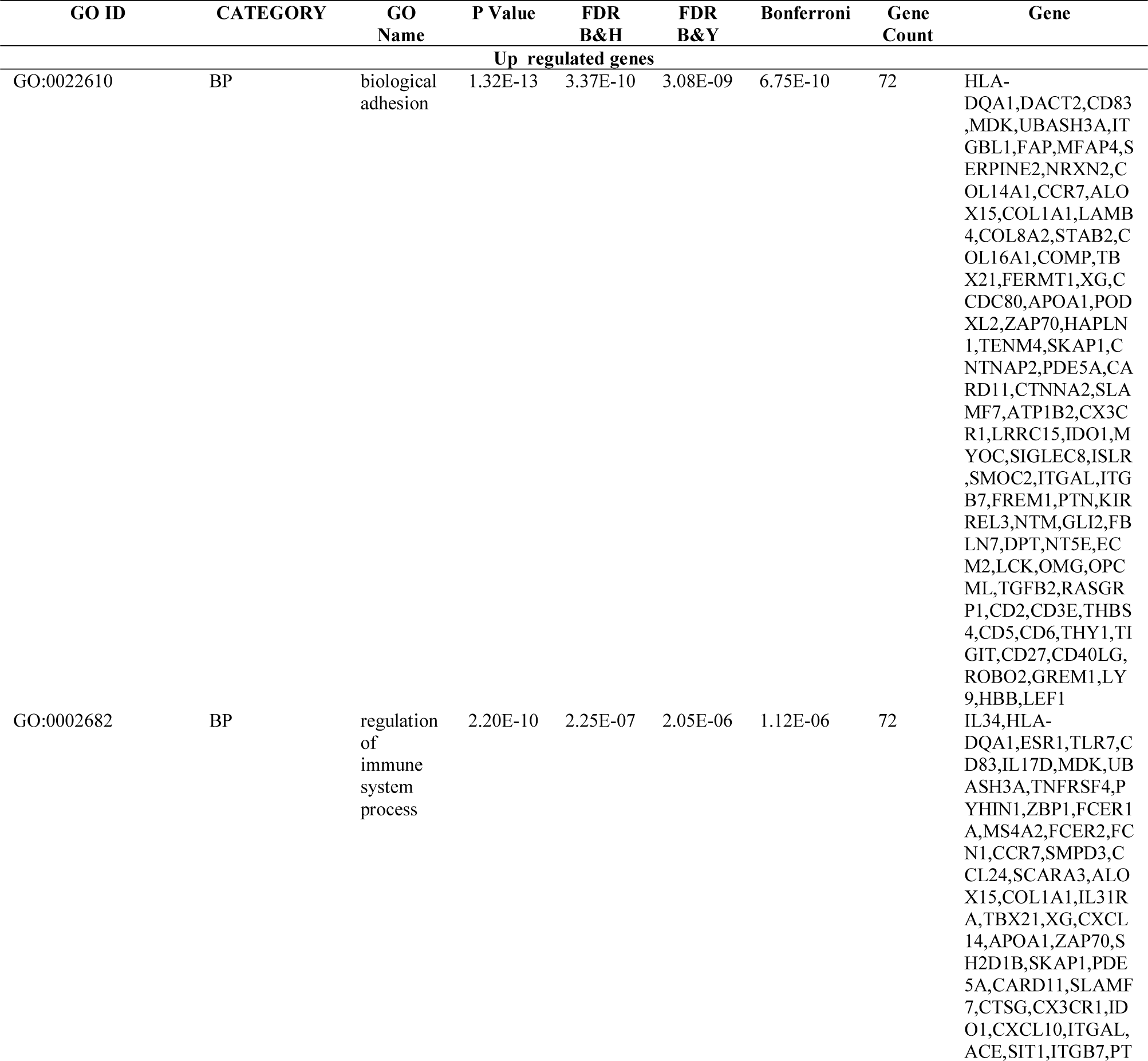
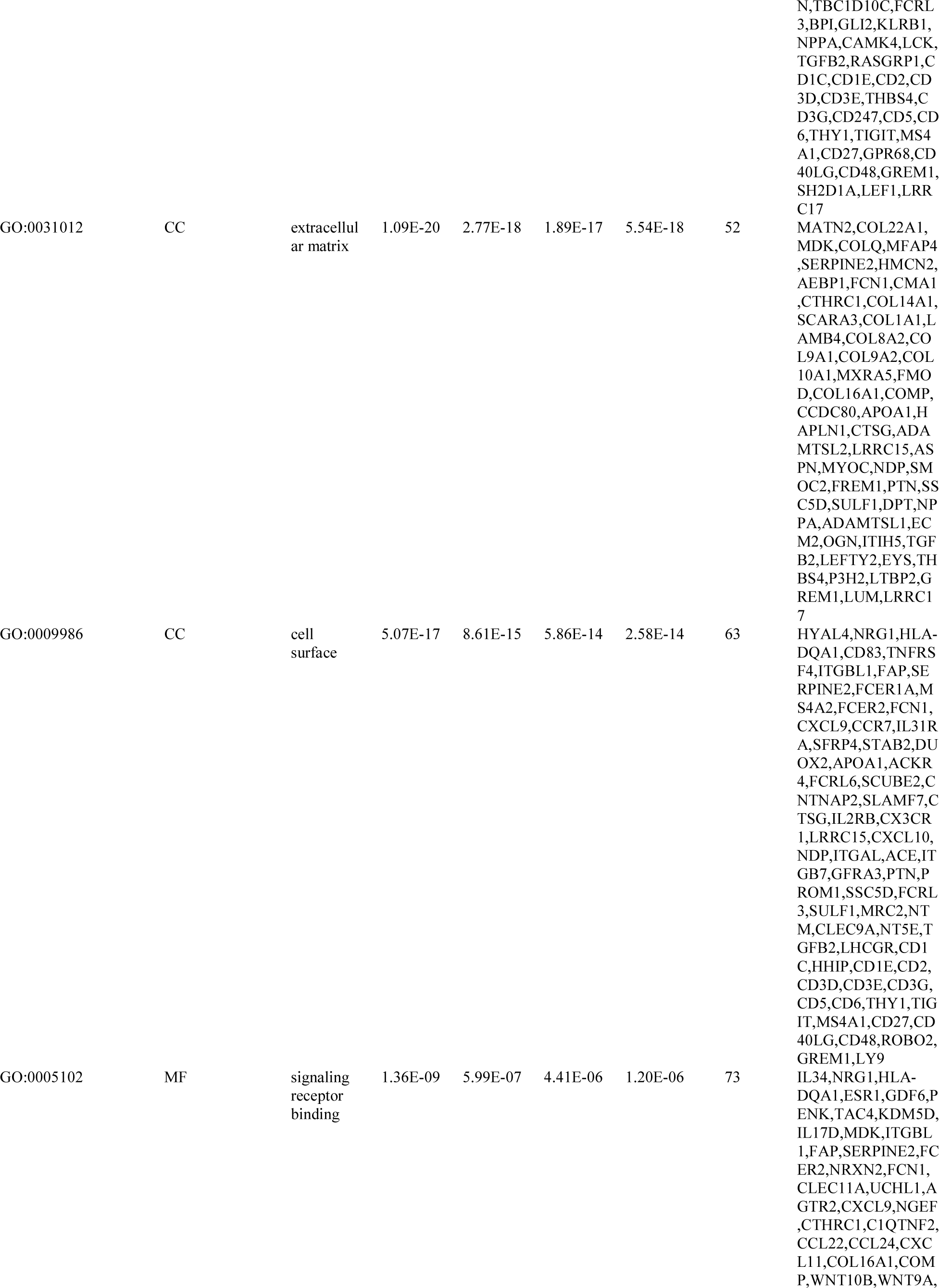
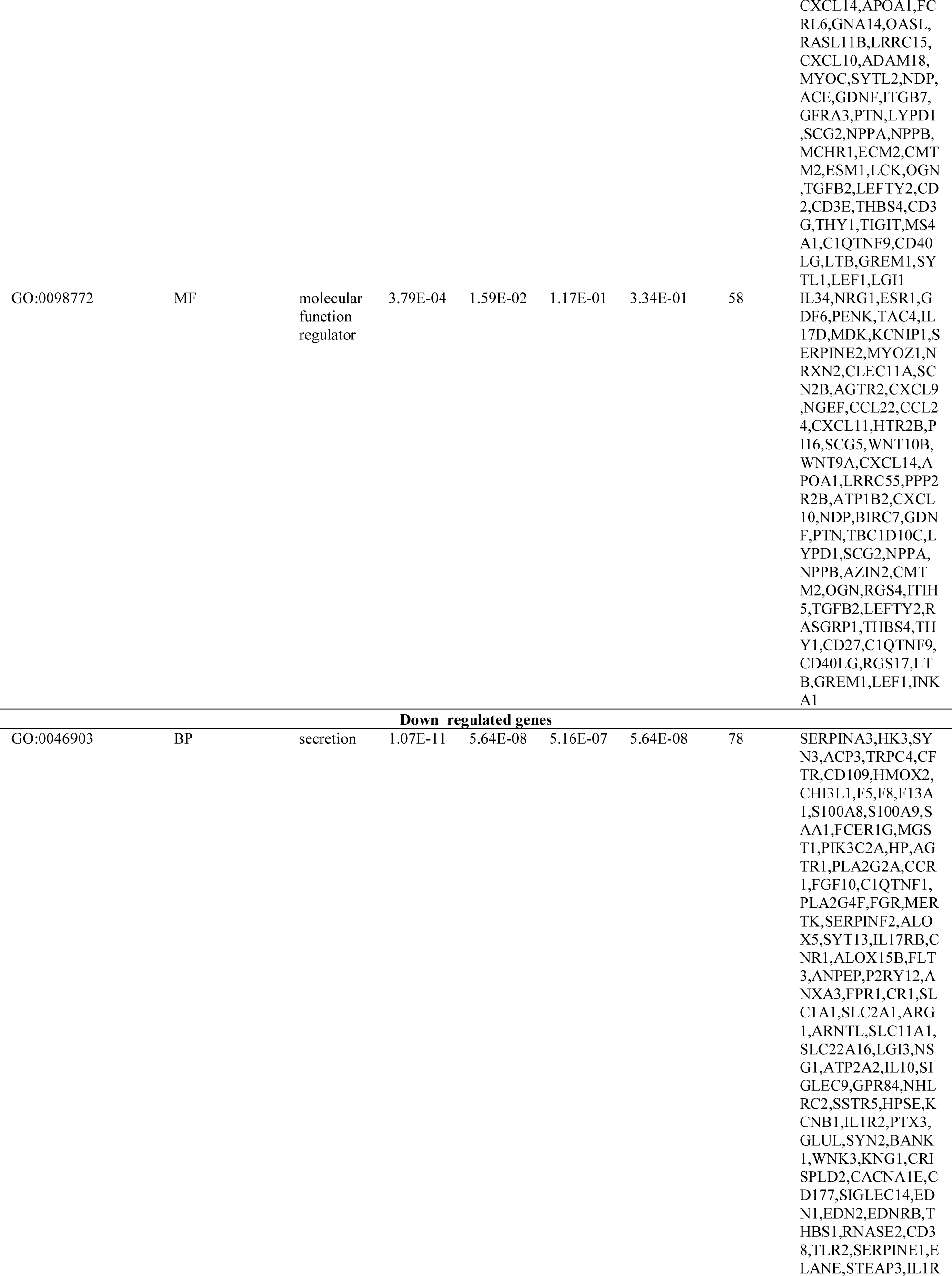
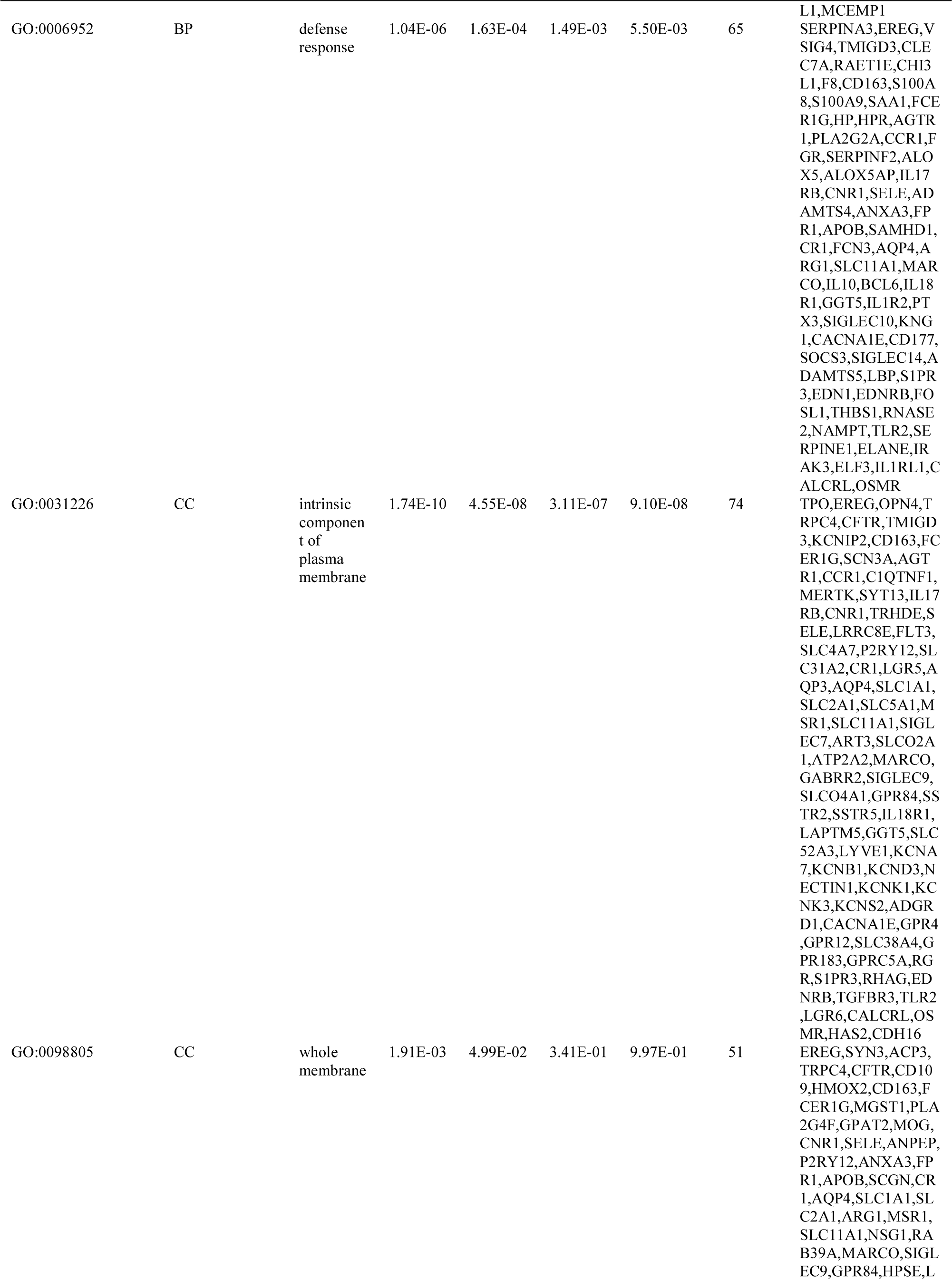
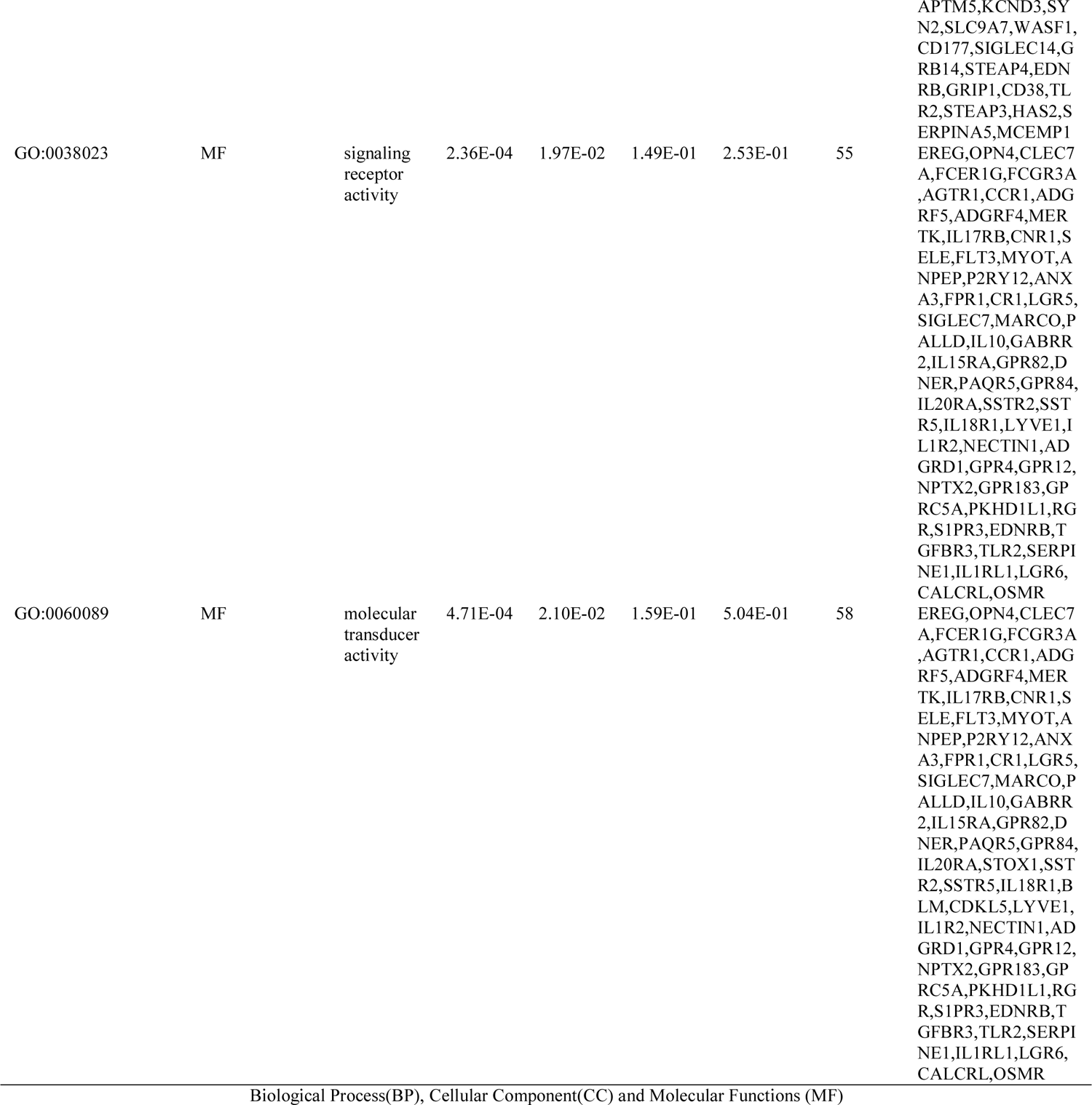
The enriched GO terms of the up and down regulated differentially expressed genes

**Table 4.**
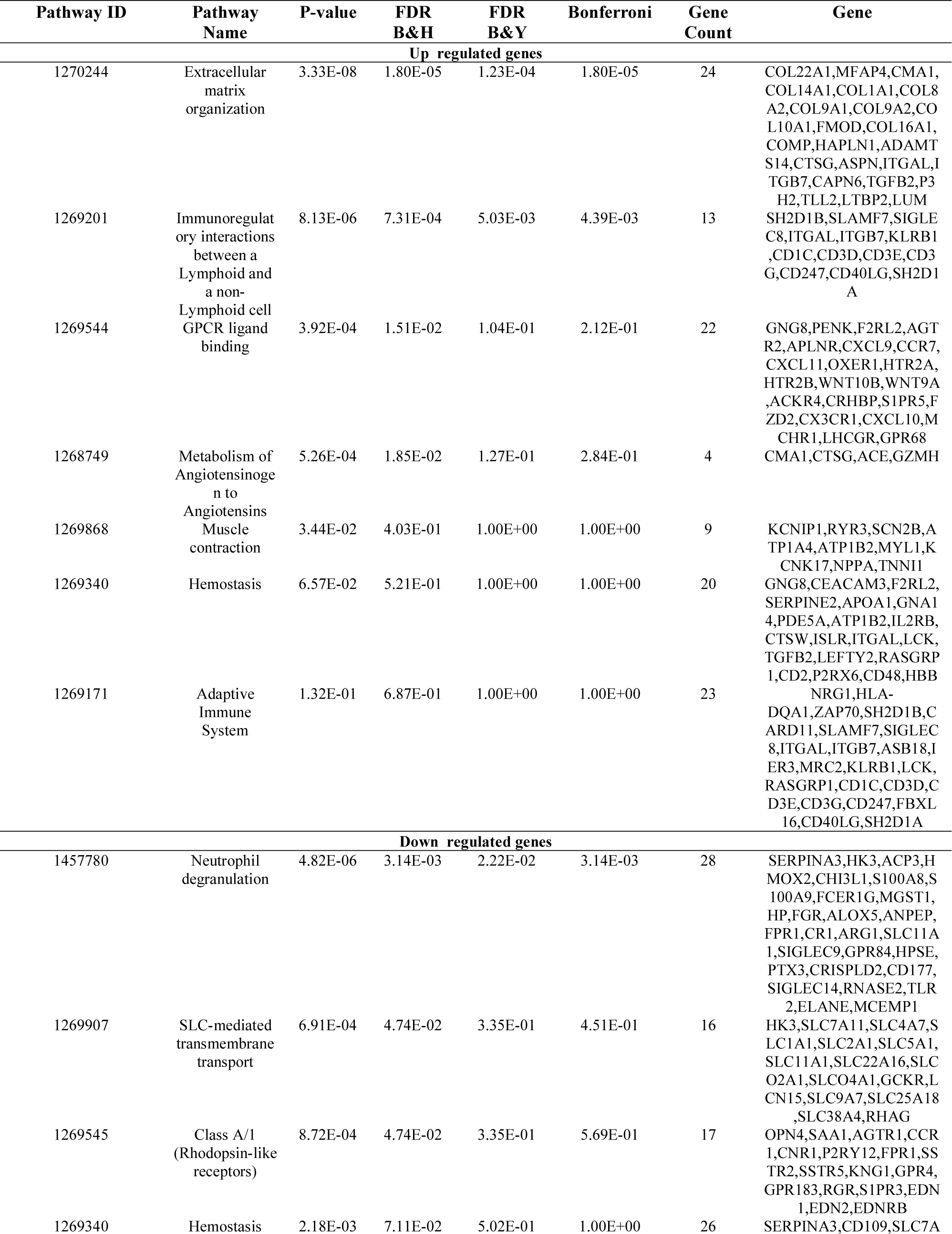
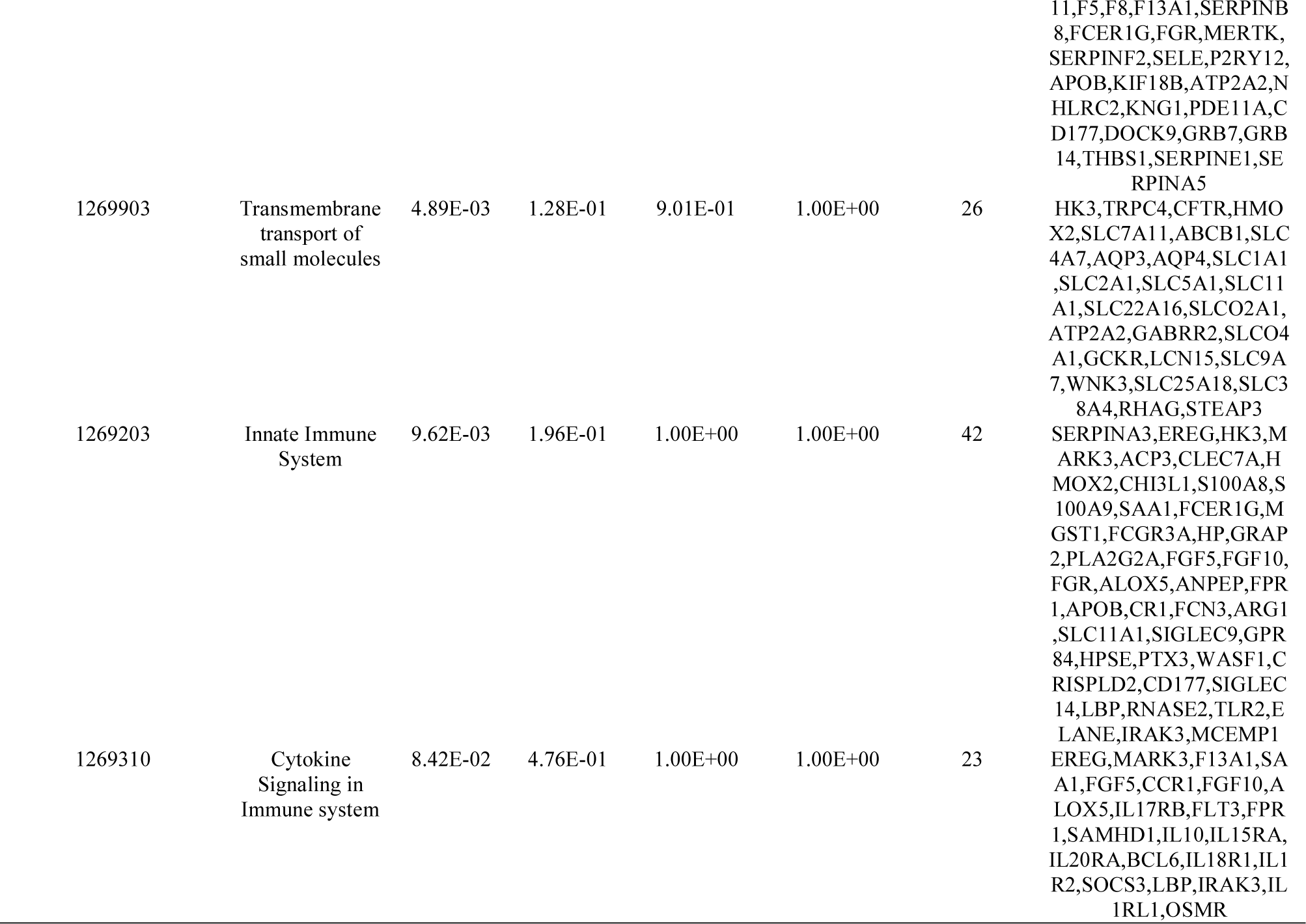
The enriched pathway terms of the up and down regulated differentially expressed genes

### Protein-protein interaction (PPI) network and module analysis

Based on the HiPPIE interactome database, the PPI network for the DEGs (including 6541 nodes and 13909 edges) was constructed (Fig. 3A). Up regulated gene with higher node degree, higher betweenness centrality, higher stress centrality and higher closeness centrality were as follows: ESR1, PYHIN1, PPP2R2B, LCK, TP63 and so on. Down regulated genes had higher node degree, higher betweenness centrality, higher stress centrality and higher closeness centrality were as follows PCLAF, CFTR, TK1, ECT2, FKBP5 and so on. The node degree, betweenness centrality, stress centrality and closeness centrality are listed in Table 5.

**Fig. 3.**
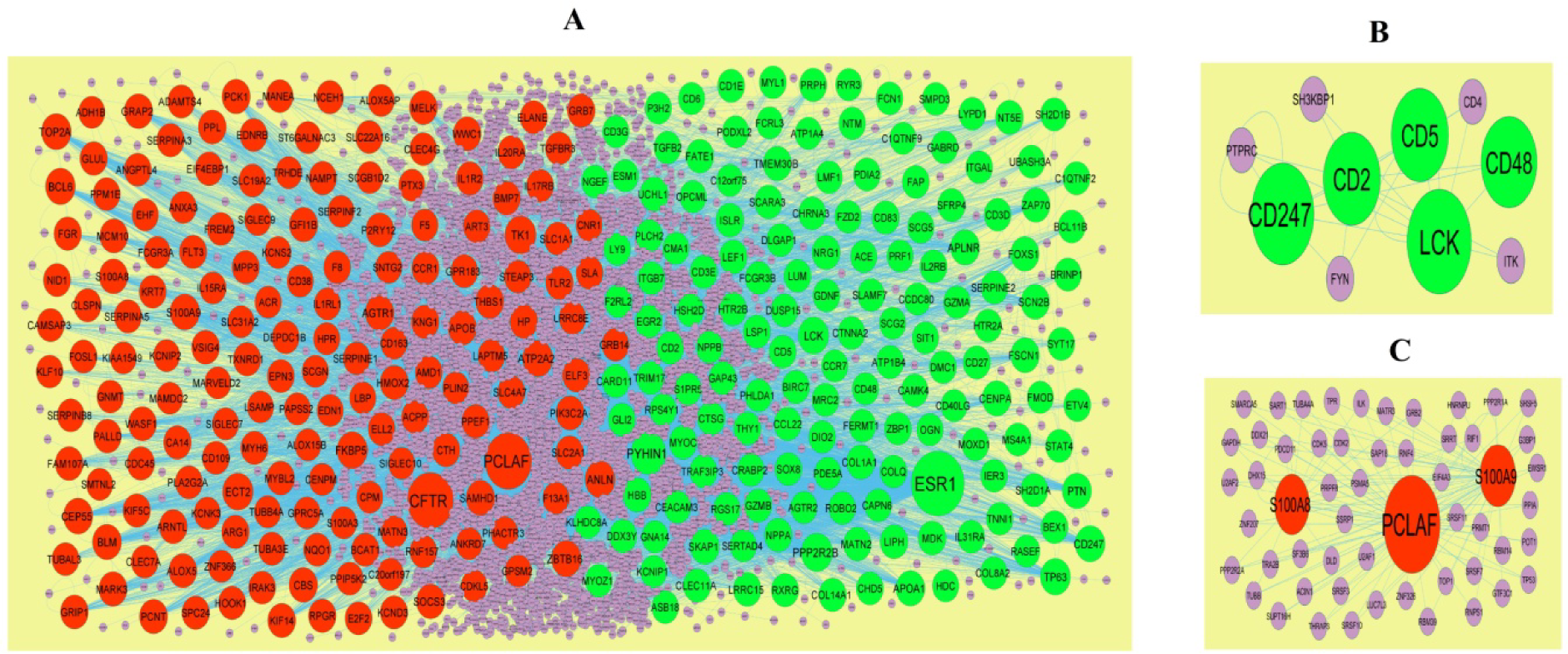
PPI network and the most significant modules of DEGs. (A) The PPI network of DEGs was constructed using Cytoscape (B) The most significant module was obtained from PPI network with 4 nodes and 6 edges for up regulated genes (C) The most significant module was obtained from PPI network with 6 nodes and 10 edges for down regulated genes. Up regulated genes are marked in green; down regulated genes are marked in red

**Table 5.**
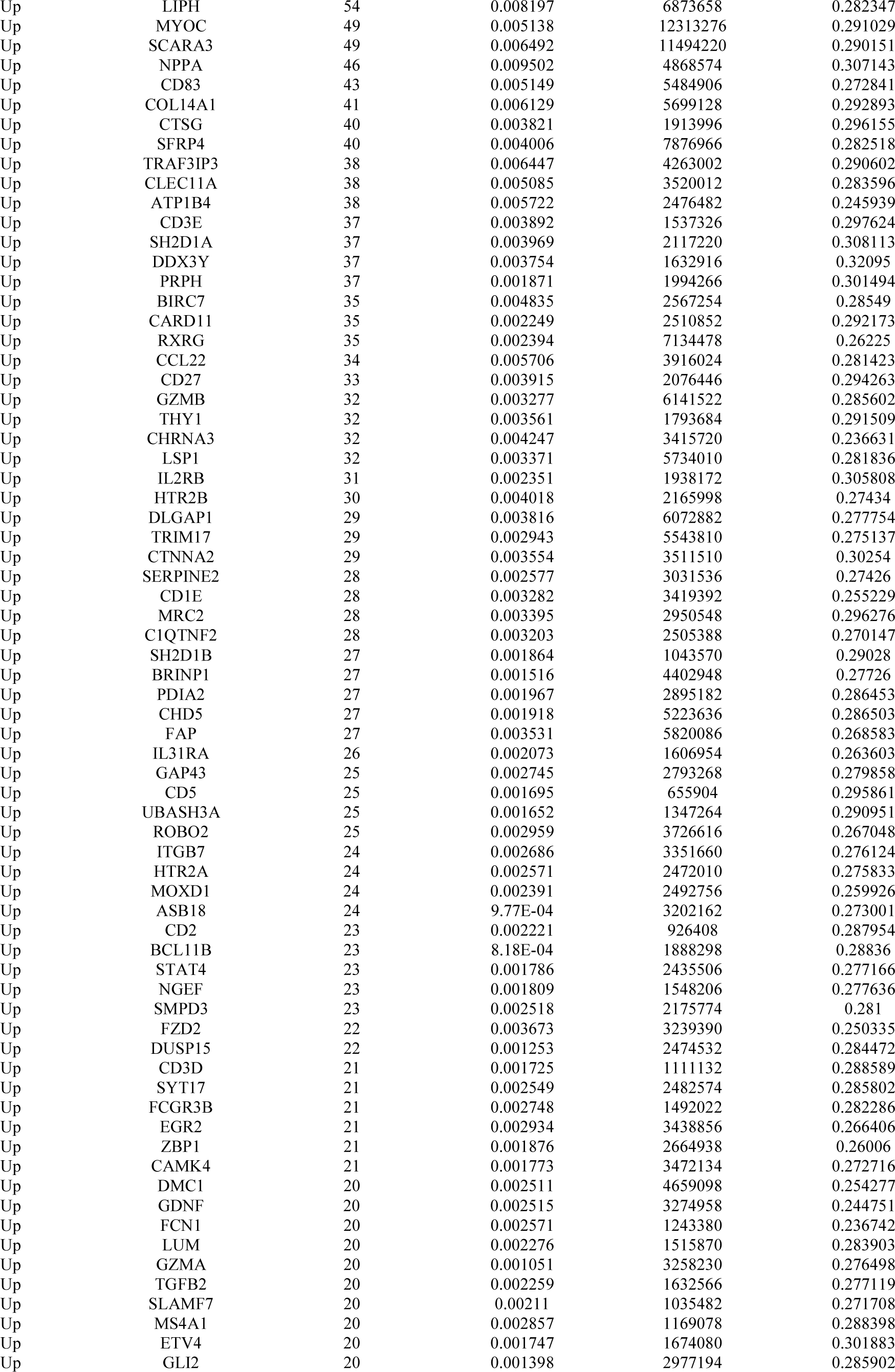
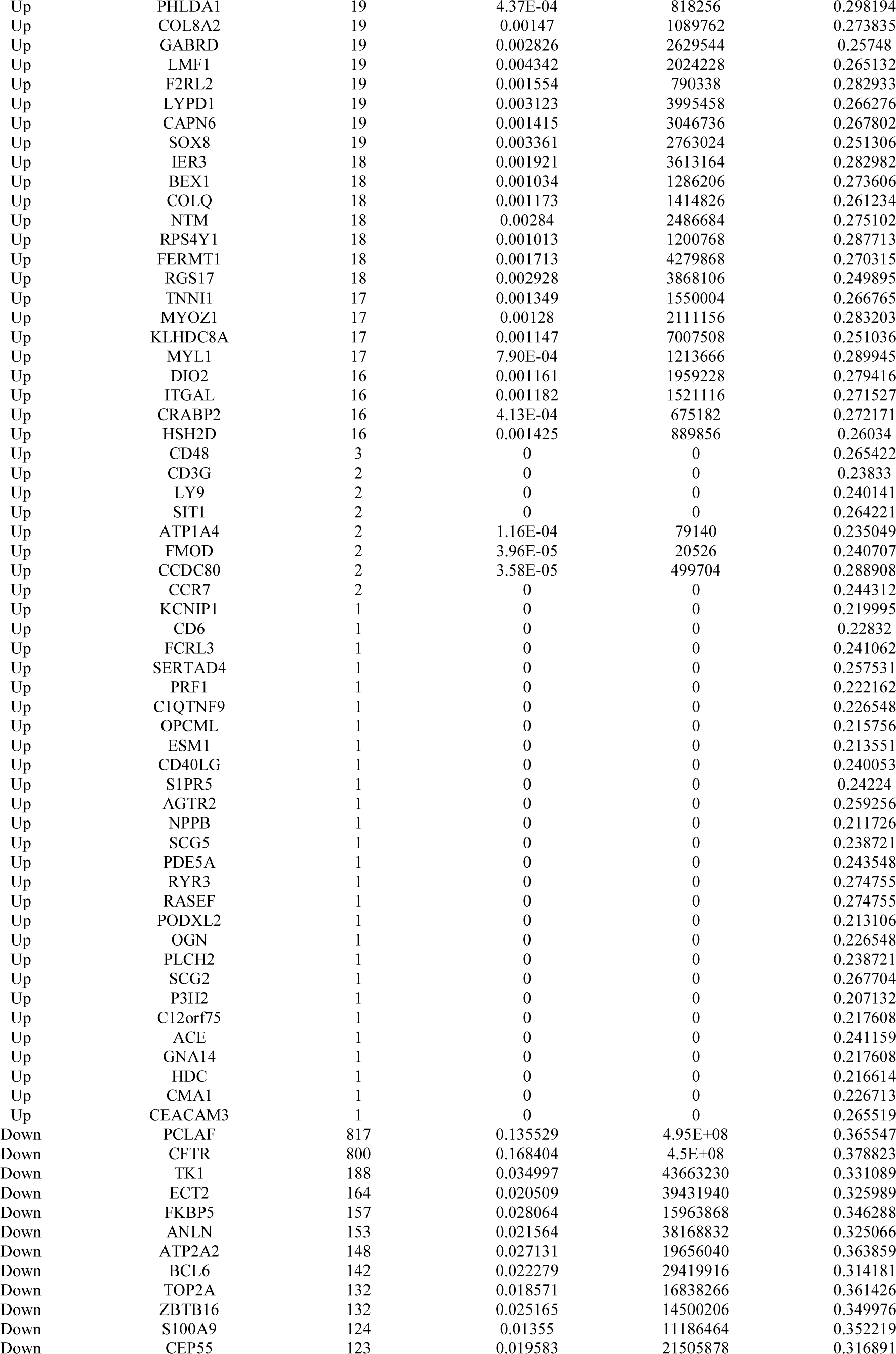
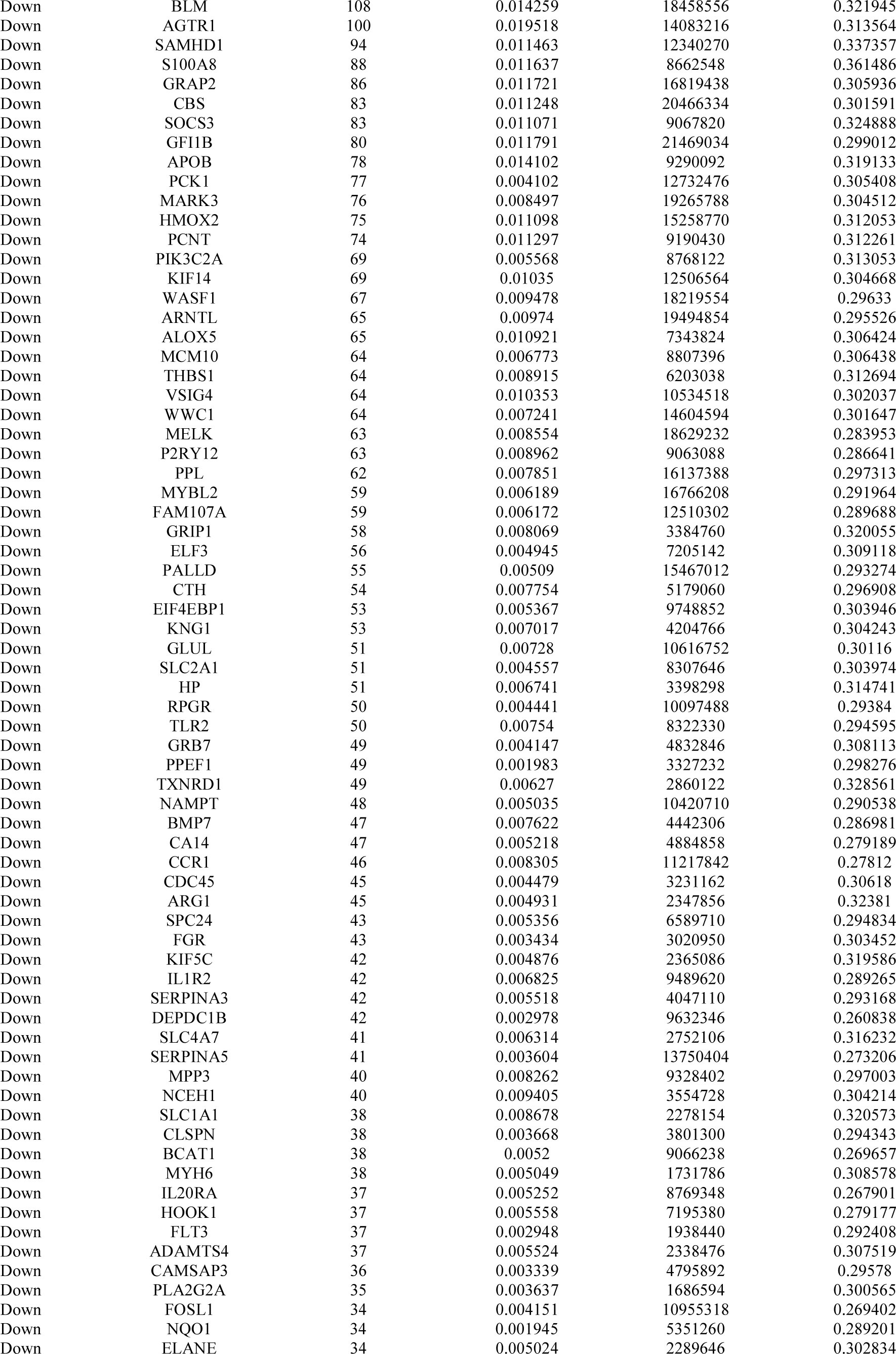
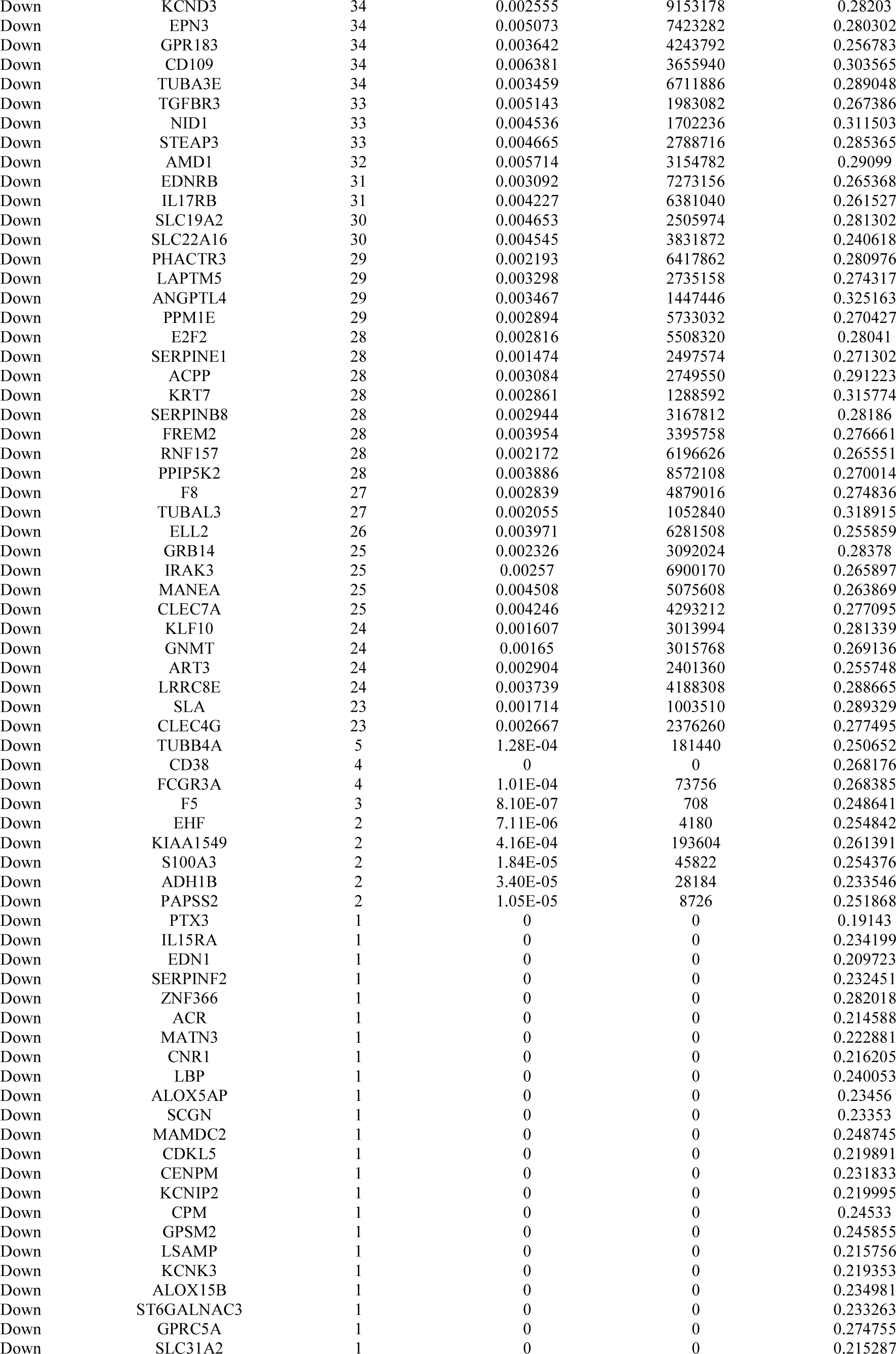
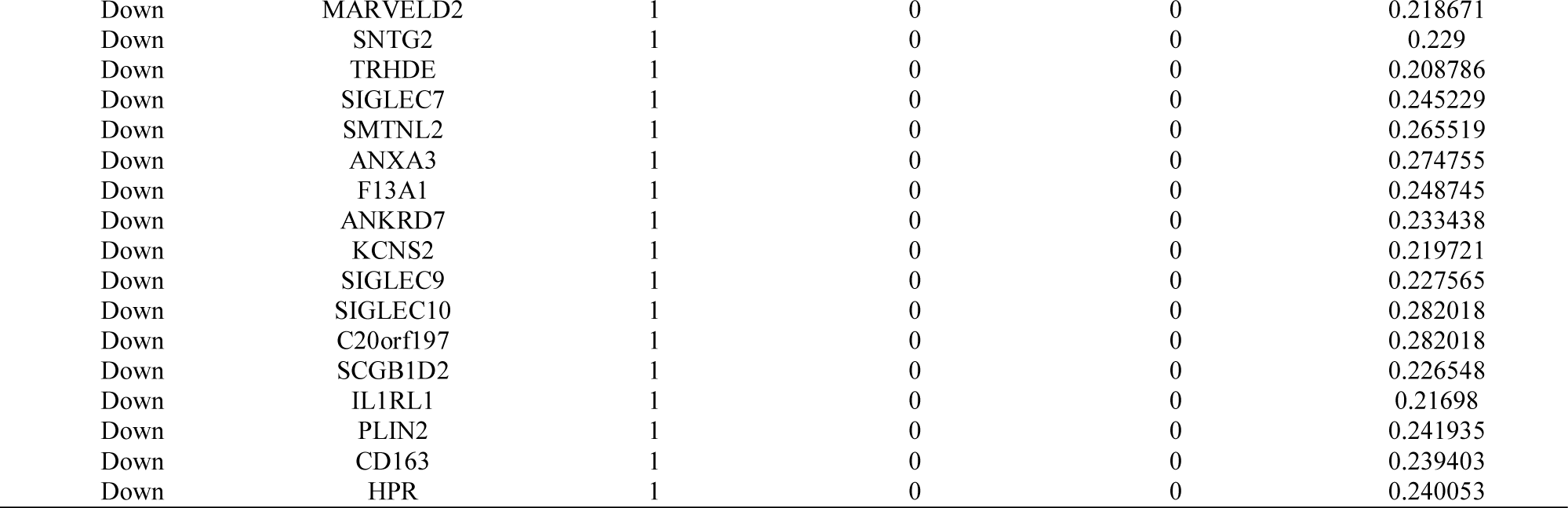
Topology table for up and down regulated genes.

Additionally, two significant modules, including module 1 (10 nodes and 24 edges) and module 2 (5 nodes and 10 edges) (Fig. 3B) and module 3 (55 nodes and 115 edges), were acquired by PEWCC1 plug-in (Fig. 3C). Furthermore, GO terms and REACTOME pathways were significantly enriched by module 1, including adaptive immune system, immunoregulatory interactions between a lymphoid and a non-lymphoid cell, hemostasis, biological adhesion and regulation of immune system process. Meanwhile, the nodes in module 2 were significantly enriched in GO terms and REACTOME pathways, including neutrophil degranulation and secretion.

### Target gene - miRNA regulatory network construction

Associations between 2063 miRNAs and their 319 target genes were collected from the target gene - miRNA regulatory network (Fig. 4). MiRNAs of hsa-mir-4533, hsa-mir-548ac, hsa-mir-548i, hsa-mir-5585-3p, hsa-mir-6750-3p, hsa-mir-200c-3p, hsa-mir-1273g-3p, hsa-mir-1244, hsa-mir-4789-3p and hsa-mir-766-3p, which exhibited a high degree of interaction, were selected from this network. Furthermore, the results also showed that FSCN1 was the target of hsa-mir-4533, ESR1 was the target of hsa-mir-548ac, TMEM30B was the target of hsa-mir-548i, SCN2B was the target of hsa-mir-5585-3p, CENPA was the target of hsa-mir-6750-3p, FKBP5 was the target of hsa-mir-200c-3p, PCLAF was the target of hsa-mir-1273g-3p, CEP55 was the target of hsa-mir-1244, ATP2A2 was the target of hsa-mir-4789-3p and TK1 was the target of hsa-mir-766-3p, and are listed in Table 6.

**Fig. 4.**
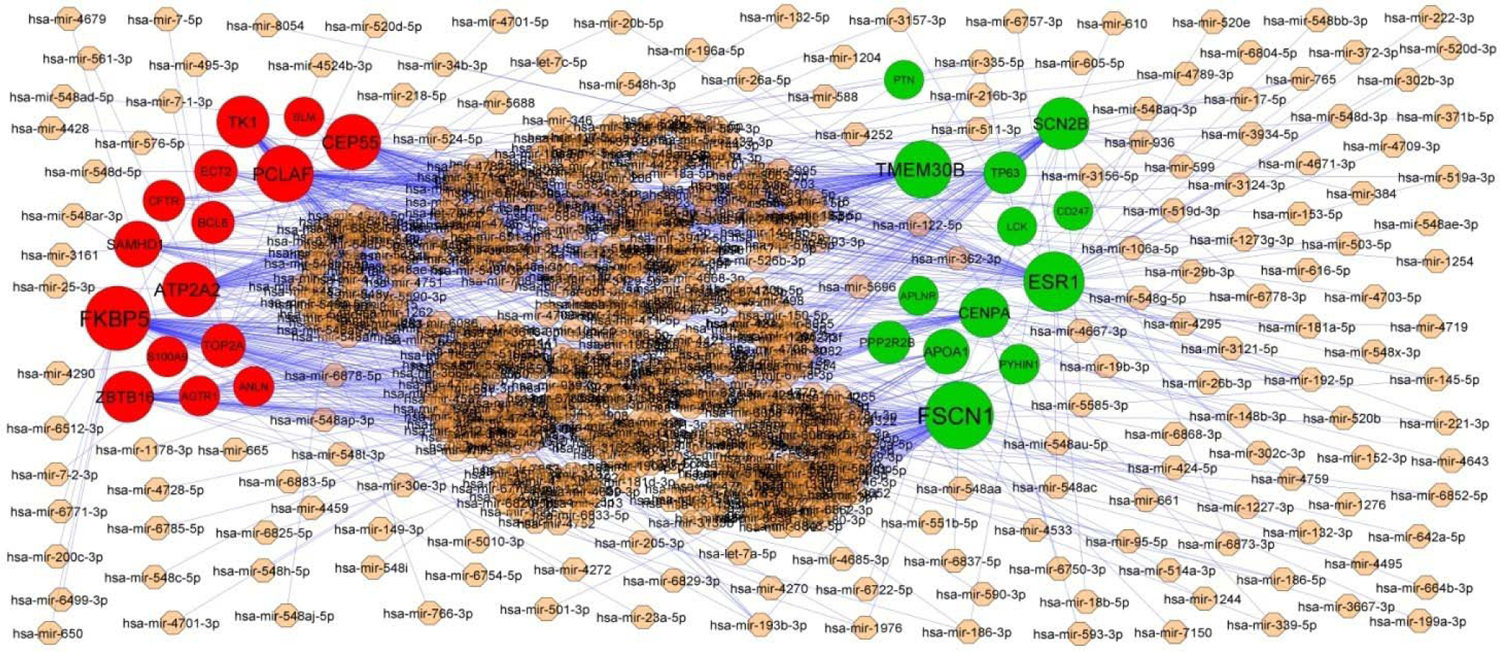
Target gene - miRNA regulatory network between target genes. The light orange color diamond nodes represent the key miRNAs; up regulated genes are marked in green; down regulated genes are marked in red.

**Table 6.**
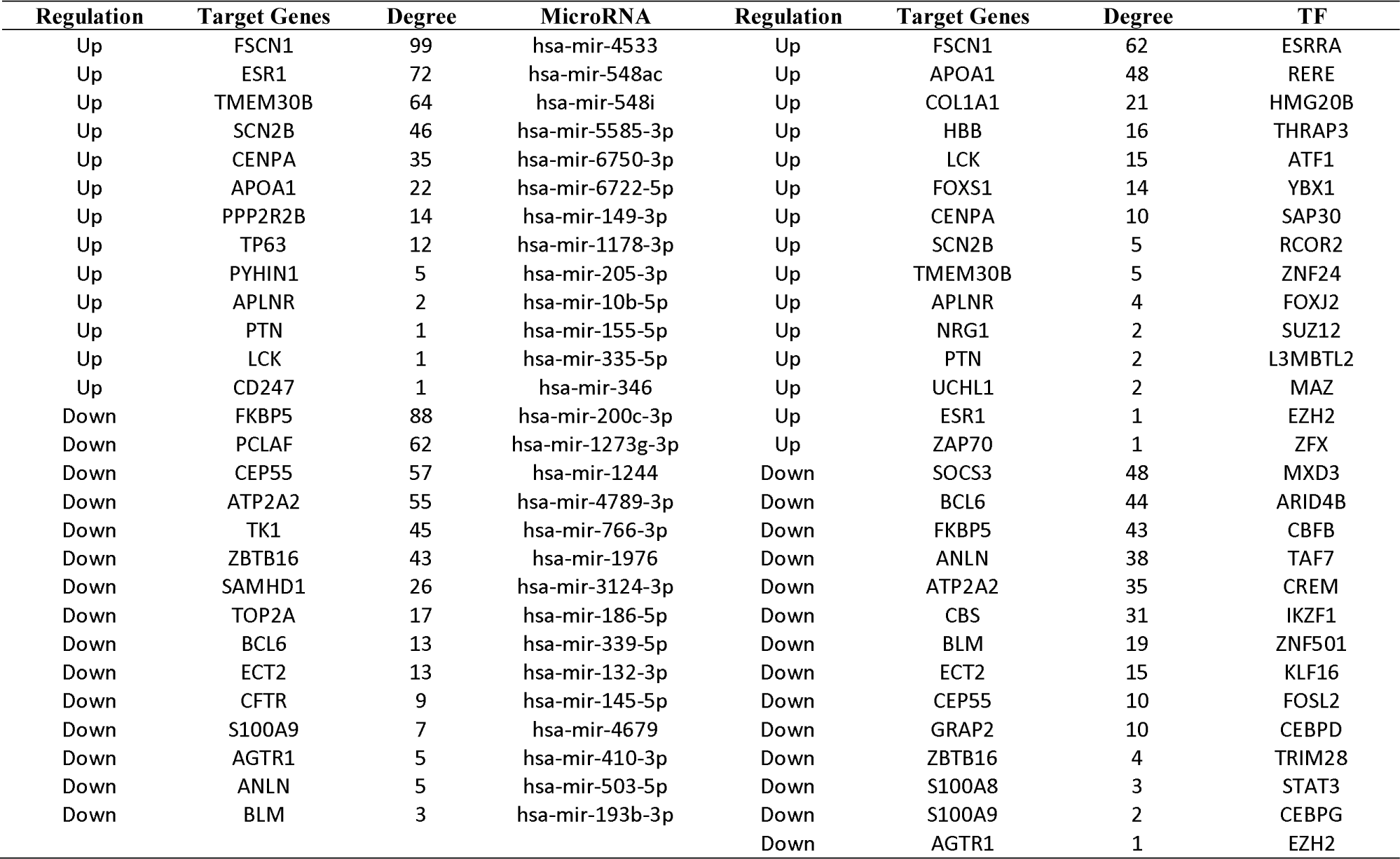
miRNA - target gene and TF - target gene interaction

### Target gene - TF regulatory network construction

Associations between 330 TFs and their 247 target genes were collected from the target gene - TF regulatory network (Fig. 5). TFs of ESRRA, RERE, HMG20B, THRAP3, ATF1, MXD3, ARID4B, CBFB, TAF7 and CREM, which exhibited a high degree of interaction, were selected from this network. Furthermore, the results also showed that FSCN1 was the target of ESRRA, APOA1 was the target of RERE, COL1A1 was the target of HMG20B, HBB was the target of THRAP3, LCK was the target of ATF1, SOCS3 was the target of MXD3, BCL6 was the target of ARID4B, FKBP5 was the target of CBFB, ANLN was the target of TAF7 and ATP2A2 was the target of CREM, and are listed in Table 6.

**Fig. 5.**
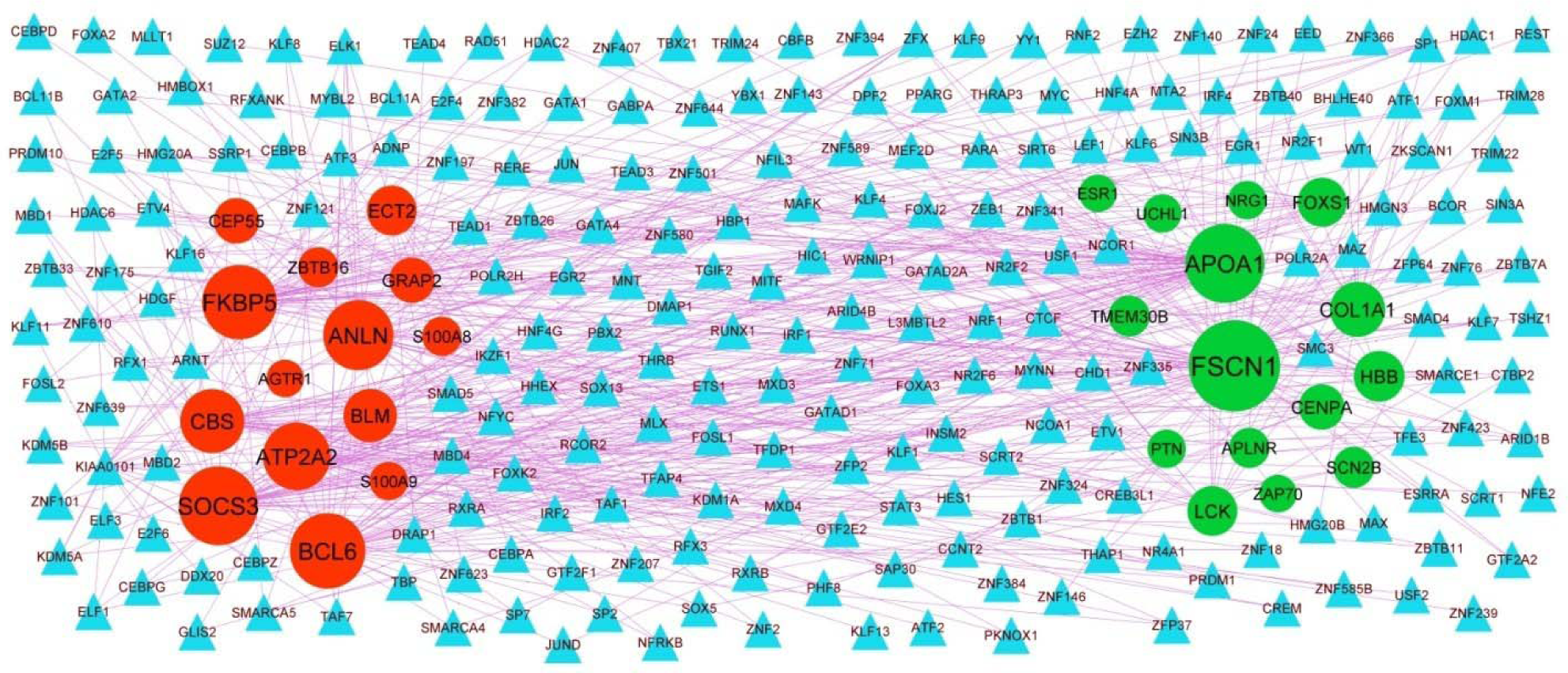
Target gene - TF regulatory network between target genes. The sky blue color triangle nodes represent the key TFs; up regulated genes are marked in green; down regulated genes are marked in red.

### Receiver operating characteristic (ROC) curve analysis

First of all, we performed the ROC curve analysis among 10 hub genes based on the GSE141910. The results showed that ESR1, PYHIN1, PPP2R2B, LCK, TP63, PCLAF, CFTR, TK1, ECT2 and FKBP5 achieved an AUC value of > 0.7, L L demonstrating that these ten genes have high sensitivity and specificity for HF, suggesting they can be served as biomarkers for the diagnosis of HF (Fig. 6).

**Fig. 6.**
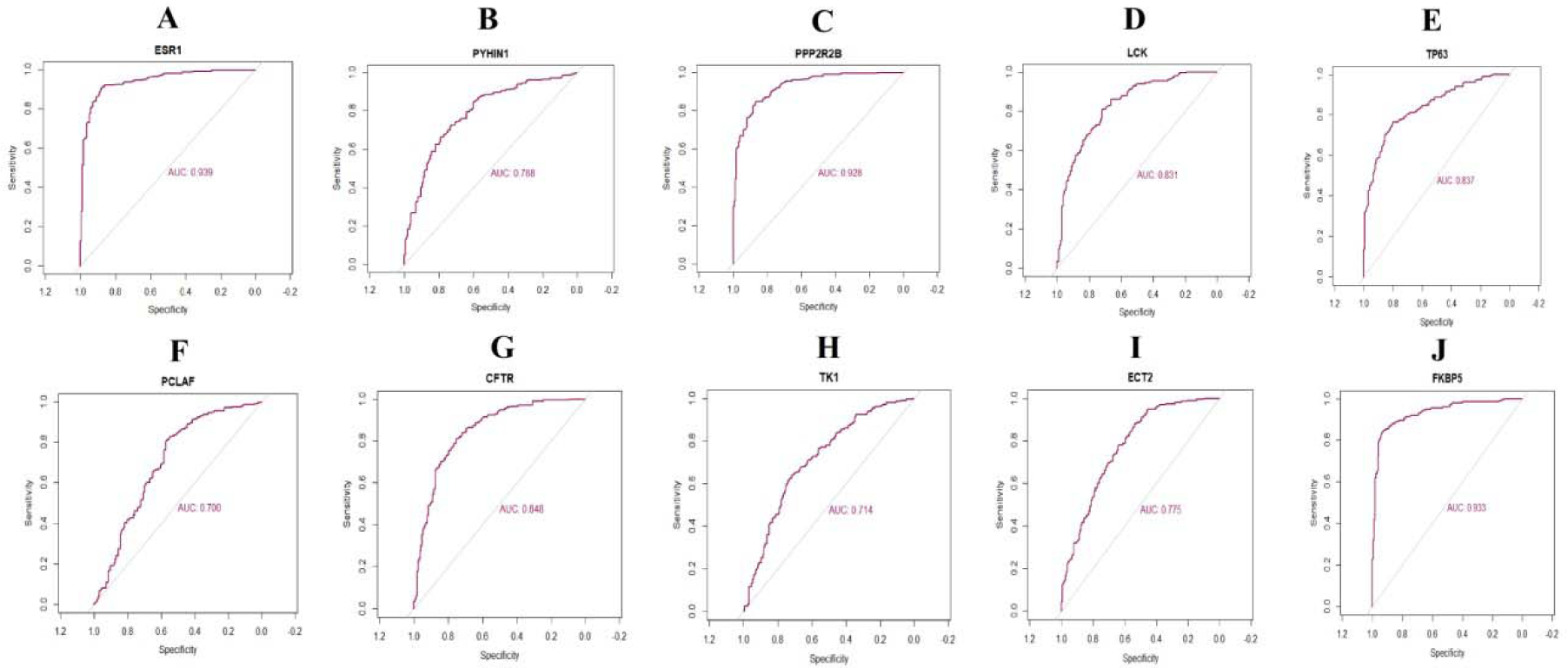
ROC curve analyses of hub genes. A) ESR1 B) PYHIN1 C) PPP2R2B D) LCK E) TP63 F) PCLAF G) CFTR H) TK1 I) ECT2 J) FKBP5

### RT-PCR analysis

RT-PCR was used to validate the hub genes between normal and HF cell lines. The results suggested that the mRNA expression level of ESR1, PYHIN1, PPP2R2B, LCK and TP63 were significantly increased in HF compared with that in normal, while PCLAF, CFTR, TK1, ECT2 and FKBP5 were significantly decreased in HF compared with that in normal and are shown in Fig. 7.

**Fig. 7.**
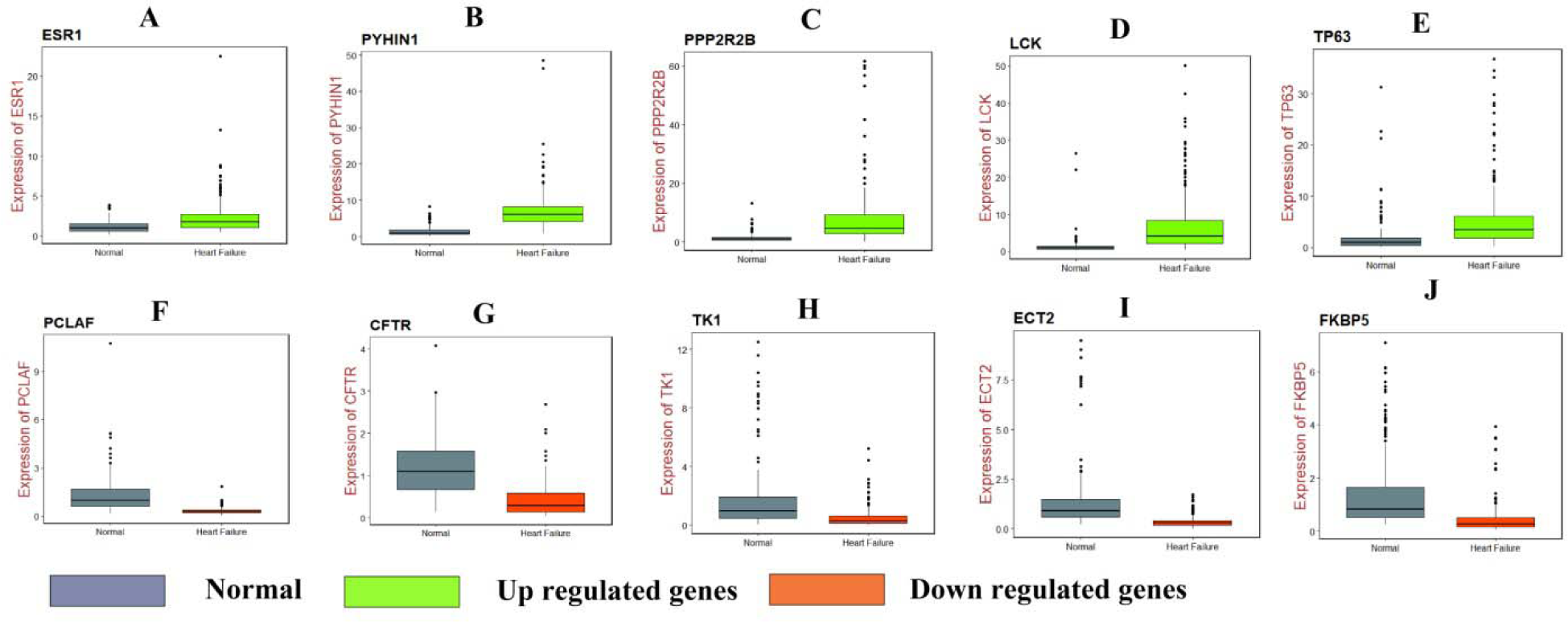
RT-PCR analyses of hub genes. A) ESR1 B) PYHIN1 C) PPP2R2B D) LCK E) TP63 F) PCLAF G) CFTR H) TK1 I) ECT2 J) FKBP5

### Identification of Candidate Small Molecules

In the present study docking simulations are performed to spot the active site and foremost interactions accountable for complex stability with the receptor binding sites. In heart failure recognized over expressed genes and their proteins of x-ray crystallographic structure are chosen from PDB for docking studies. Most generally, medications containing benzothiadiazine ring hydrochlorothiazide are used in heart failure either alone or in conjunction with other drugs, based on this the molecules containing heterocyclic ring of benzothiadiazine are designed and hydrochlorothiazide is uses as a reference standard. Docking experiments using Sybyl-X 2.1.1. drug design perpetual software were used on the designed molecules. Docking studies were performed in order to understand the biding interaction of standard hydrochlorothiazide and designed molecules on over expressed protein. The X-RAY crystallographic structure of one proteins from each over expressed genes of ESR1, LCK, PPP2R2B, PYHIN1, TP63 and their co-crystallised protein of PDB code 4PXM, 1KSW, 2HV7, 3VD8 and 6RU6 respectively were selected for the docking studies to identify and predict the potential molecule based on the binding score with the protein and successful in heart failure. For the docking tests, a total of 34 molecules were built and the molecule with binding score greater than 5 is believed to be good. The designed molecules obtained docking score of 5 to 7 were HIM10, HTZ5, HIM6, HTZ31, HIM3, HIM14, HIM1, HIM7 and HIM11, HIM16, HTZ9, HIM17, HIM12, HTZ12, HIM6, HTZ7, HIM10, HTZ3 and HIM8, HTZ9, HIM6, HIM4, HIM13, HTZ16, HIM9, HIM7, HTZ5, HIM16, HTZ7, HIM10, HIM5, HIM12, HIM15, HTZ12, HIM3, HIM14 and HIM14, HIM6, HIM17, HTZ7, HIM10, HIM1, HTZ9, HIM3, HIM16, HIM15, HIM8, HIM9, HIM7, HTZ10, HTZ3, HTZ5, HTZ1, HIM13, HTZ4, HIM11, HTZ12, HTZ14, HIM2 and HIM7, HTZ13, HTZ5, HIM15, HIM12, HIM6, HTZ11, HIM14, HTZ9, HIM11, HIM13, HIM9, HIM8, HIM10, HIM1, HIM5, HIM4, HTZ12, HIM2, HIM17, HIM3, HTZ1, HTZ8, HIM3, HTZ14, HTZ3 with proteins 4PXM and, 1KSW and 2HV7 and 3VD8 and 6RUR respectively (Fig. 8). The molecules obtained binding score of less than 5 were HTZ13, HTZ12, HTZ10, HIM3, HIM15, HIM16, HIM13, HIM8, HTZ16, HIM2, HIM4, HIM17, HTZ17, HIM11, HTZ5, HTZ3, HIM9, HTZ15, HTZ5, HTZ9, HTZ11, HIM5, HTZ8 and HTZ14, HIM14, HTZ13, HIM13, HTZ16, HIM2, HIM3, HTZ10, HIM7, HIM1, HTZ1, HTZ4, HIM8, HIM5, HTZ2, HIM9, HTZ5, HTZ15, HTZ3, HIM4, HIM15, HTZ17, HTZ8, HTZ11 and HTZ14, HIM2, HIM1, HTZ11, HIM17, HTZ13, HTZ4, HTZ2, HIM3, HTZ15, HTZ8, HTZ17, HTZ1, HTZ3 and HTZ8, HIM4, HTZ16, HTZ15, HIM5, HTZ11, HTZ13, HIM3, HTZ17, HTZ2 and HTZ7, HTZ4, HTZ2, HTZ17, HTZ15 with proteins 4PXM and, 1KSW and 2HV7 and 3VD8 and 6RUR respectively. The molecules obtained very less binding score are HTZ1, HIM12, HTZ2, HTZ4 with protein 4PXM and the standard hydrochlorothiazide (HTZ) obtained less binding score with all proteins, the values are depicted in Table 7.

**Fig. 8.**
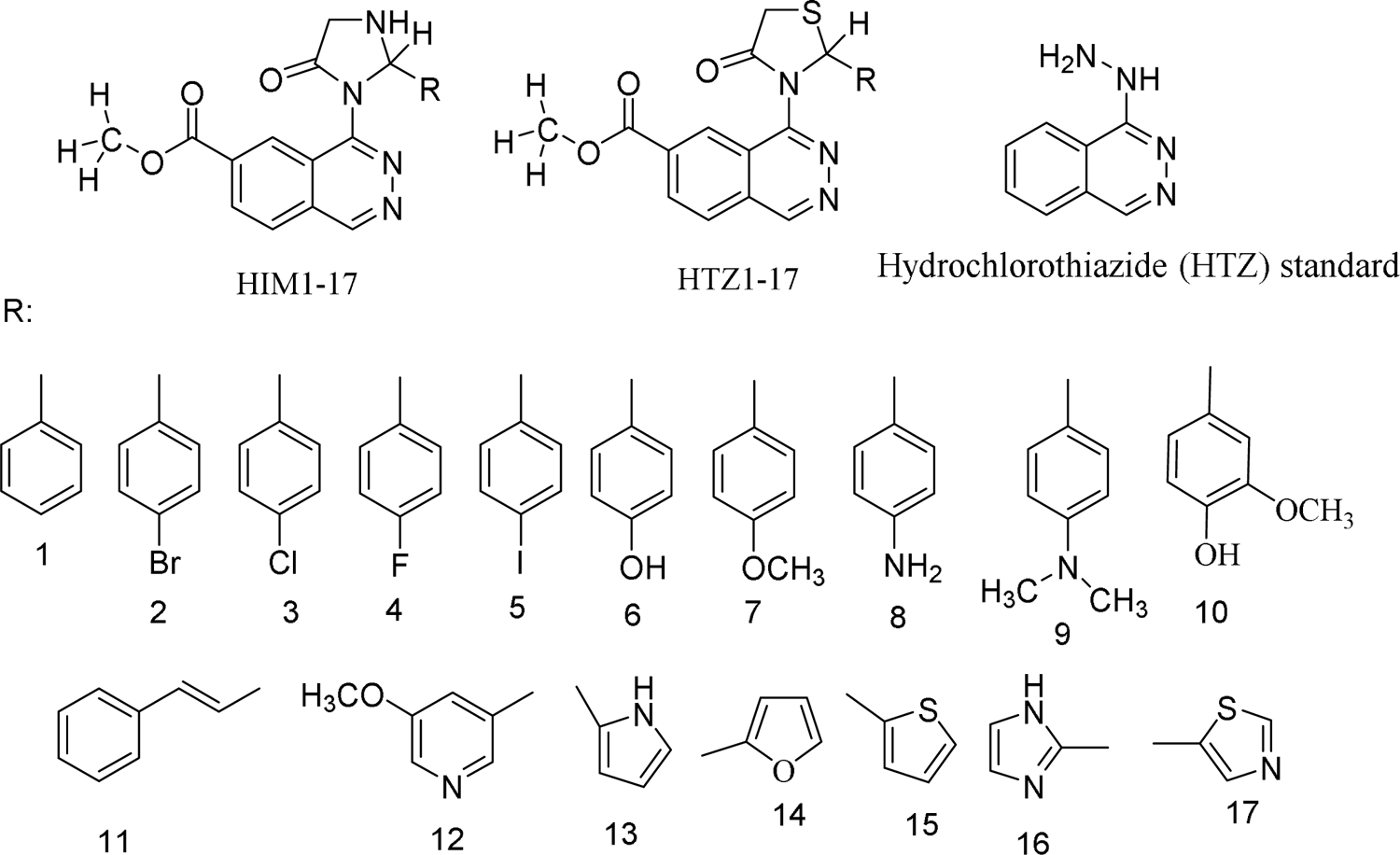
Structures of Designed Molecules

**Table 7.**
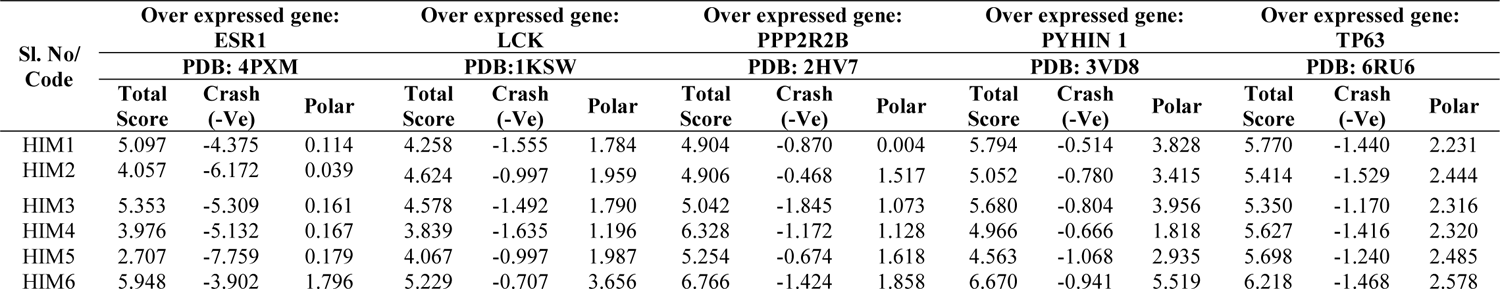
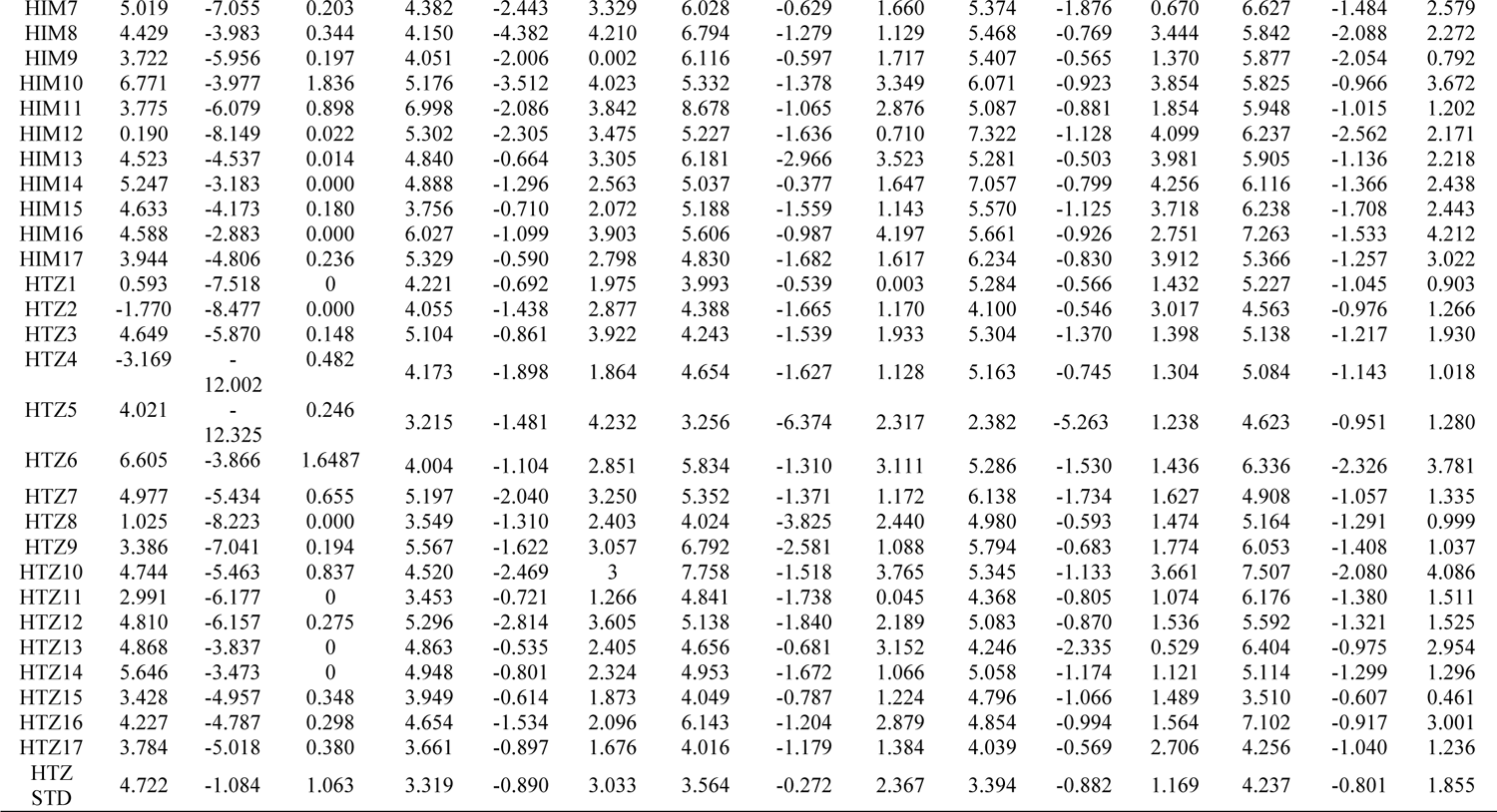
Docking results of Designed Molecules on Over Expressed Proteins

## Discussion

HF is the most prevalent form of cardiovascular disease among the elderly. A complete studies of HF, comprising pathogenic factors, pathological processes, clinical manifestations, early clinical diagnosis, clinical prevention, and drug therapy targets urgency to be consistently analyzed. In the present investigation, bioinformatics analysis was engaged to explore HF biomarkers and the pathological processes in myocardial tissues, acquired from HF groups and non heart failure groups. We analyzed GSE141910 expression profiling by high throughput sequencing obtained 881 different genes between HF groups and non heart failure groups, 442 up regulated and 439 down regulated genes. HBA2 and HBA1 have a key role in hypertension [32], but these genes might be linked with development HF. SFRP4 was linked with progression of myocardial ischemia [33]. Emmens et al. [34] and Broch et al. [35] found that PENK (proenkephalin) and IL1RL1 were up regulated in HF. ALOX15B has lipid accumulation and inflammation activity and is highly expressed in atherosclerosis [36]. Studies have shown that expression of MYH6 was associated with hypertrophic cardiomyopathy [37].

In functional enrichment analysis, some genes involved with regulation of cardiovascular system processes were enriched in HF. Liu et al. [38], Kosugi et al. [39], McMacken et al. [40], Pan and Zhang [41], Li et al. [42] and Jiang et al. [43] presented that expression of HLA-DQA1, KDM5D, UCHL1, SAA1, ARG1 and LYVE1 were associated with progression of cardiomyopathy. Hou et al. [44] and Olesen et al. [45] demonstrated that DACT2 and KCND3 were found to be substantially related to atrial fibrillation. Ge and Concannon [46], Ferjeni et al. [47], Anquetil et al. [48], Glawe et al. [49], Kawabata et al. [50], Li et al. [51], Buraczynska et al. [52], Amini et al. [53], Yang et al. [54], Du Toit et al. [55], Hirose et al. [56], Zhang et al. [57], Griffin et al. [58], Zouidi et al. [59], Trombetta et al. [60], Alharbi et al. [61], Ikarashi et al. [62], Dharmadhikari et al. [63], Sutton et al. [64] and Deng et al. [65] reported that UBASH3A, ZAP70, IDO1, ITGAL (integrin subunit alpha L). ITGB7, RASGRP1, CNR1, SLC2A1, SLC11A1, GPR84, SSTR5, KCNB1, GLUL (glutamate-ammonia ligase), BANK1, CACNA1E, LGR5, AQP3, SIGLEC7, SSTR2 and DNER (delta/notch like EGF repeat containing) could be an index for diabetes, but these genes might be responsible for progression of HF. Experiments show that expression of FAP (fibroblast activation protein alpha) [66], THBS4 [67], CD27 [68], LEF1 [69], CTHRC1 [70], ESR1 [71], CXCL9 [72], SERPINA3 [73], TRPC4 [74], F13A1 [75], PIK3C2A [76], KCNIP2 [77] and GPR4 [78] contributed to myocardial infarction. MFAP4 [79], ALOX15 [80], COL1A1 [81], APOA1 [82], PDE5A [83], CX3CR1 [84], THY1 [85], GREM1 [86], FMOD (fibromodulin) [87], NPPA (natriuretic peptide A) [88], LTBP2 [89], LUM (lumican) [90], IL34 [91], NRG1 [92], CXCL14 [93], CXCL10 [94], ACE (angiotensin I converting enzyme) [95], CFTR (ystic fibrosis transmembrane conductance regulator) [96], S100A8 [97], S100A9 [97], HP (haptoglobin) [98], AGTR1 [99], ATP2A2 [100], IL10 [101], EDN1 [102], TLR2 [103], MCEMP1 [104], TPO (thyroid peroxidase) [105], CD163 [106], IL18R1 [107], KCNA7 [108] and CALCRL (calcitonin receptor like receptor) [109] have an important role in HF. Li et al [110], Deckx et al [111], Ichihara et al [112] and Paik et al [113] showed that the SERPINE2, OGN (osteoglycin), AGTR2 and WNT10B promoted cardiac interstitial fibrosis. Cai et al [114], Mo et al [115], Sun et al [116], Martinelli et al [117], Zhao et al [118], Assimes et al [119] and Piechota et al [120] showed that CCR7, FCN1, ESM1, F8 (coagulation factor VIII), C1QTNF1, ALOX5 and MSR1 were an important target gene for coronary artery disease. STAB2 have been suggested to be associated with venous thromboembolic disease [121]. Genes such as COMP (cartilage oligomeric matrix protein) [122], CHI3L1 [123], PLA2G2A [124], P2RY12 [125], CR1 [126], HPSE (heparanase) [127], PTX3 [128] and SERPINE1 [129] were related to atherosclerosis. CCDC80 [130], CMA1 [131], MDK (midkine) [132], GNA14 [133], SCG2 [134], NPPB (natriuretic peptide B) [135], FGF10 [136], ARNTL (aryl hydrocarbon receptor nuclear translocator like) [137], WNK3 [138], EDNRB (endothelin receptor type B) [139], THBS1 [140], SELE (selectin E) [141], SLC4A7 [142], AQP4 [143] and KCNK3 [144] are thought to be responsible for progression of hypertension, but these genes might to be associated with progression of HF. CNTNAP2 [145], GLI2 [146], DPT (dermatopontin) [147], AEBP1 [148], ITIH5 [149], CXCL11 [150], GDNF (glial cell derived neurotrophic factor) [151], MCHR1 [152], FLT3 [153], ELANE (elastase, neutrophil expressed) [154], OSMR (oncostatin M receptor) [155] and IL15RA [156] are involved in development of obesity, but these genes might be key for progression of HF. CTSG (cathepsin G) is a protein coding gene plays important roles in aortic aneurysms [157]. Evidence from Safa et al. [158], Chen et al. [159], Zhou et al. [160], Hu et al. [161], Lou et al. [162], Zhang et al. [163] and Chen et al. [164] study indicated that the expression of CCL22, CCR1, FPR1, KNG1, CRISPLD2, CD38 and GPRC5A were linked with progression of ischemic heart disease. Li et al. [165] showed that STEAP3 expression can be associated with cardiac hypertrophy progression.

The HiPPIE interactome database was used to construct the PPI network, and modules analysis was performed. We finally screened out up regulated hub genes and down regulated hub genes, including ESR1, PYHIN1, PPP2R2B, LCK, TP63, PCLAF, CD247, CD2, CD5, CD48, CFTR, TK1, ECT2, FKBP5, S100A9 and S100A8 from the PPI network and its modules. TP63 might serve as a potential prognostic factor in cardiomyopathy [166]. The expression of FKBP5 is related to the progression of coronary artery disease [167]. CD247 plays a central role in hypertension [168], but this gene might be involved in the HF. PYHIN1, PPP2R2B, LCK (LCK proto-oncogene, Src family tyrosine kinase), PCLAF (PCNA clamp associated factor), TK1, ECT2, CD2, CD5 and CD48 might be the novel biomarker for HF.

The miRNet database and NetworkAnalyst database were used to construct the target gene - miRNA regulatory network and target gene - TF regulatory network. We finally screened out target genes, miRNA, TFs, including FSCN1, ESR1, TMEM30B, SCN2B, CENPA, FKBP5, PCLAF, CEP55, ATP2A2, TK1, APOA1, COL1A1, HBB, LCK, SOCS3, BCL6, ANLN, hsa-mir-4533, hsa-mir-548ac, hsa-mir-548i, hsa-mir-5585-3p, hsa-mir-6750-3p, hsa-mir-200c-3p, hsa-mir-1273g-3p, hsa-mir-1244, hsa-mir-4789-3p, hsa-mir-766-3p, ESRRA, RERE, HMG20B, THRAP3, ATF1, MXD3, ARID4B, CBFB, TAF7 and CREM from the target gene - miRNA regulatory network and target gene - TF regulatory network. SCN2B [169] and SOCS3 [170] are considered as a markers for HF and might be a new therapeutic target. BCL6 levels are correlated with disease severity in patients with atherosclerosis [171]. A previous study showed that hsa-mir-1273g-3p [172], hsa-mir-4789-3p [173] and ATF1 [174] could involved in hypertension, but these markers might be responsible for progression of HF. hsa-miR-518f, was demonstrated to be associated with cardiomyopathy [175]. An evidence demonstrating a role for ESRRA (estrogen related receptor alpha) [176] and THRAP3 [177] in diabetes, but these genes might be liable for development of HF. FSCN1, TMEM30B, CENPA (centromere protein A), CEP55, HBB (hemoglobin subunit beta), ANLN (anillin actin binding protein), hsa-mir-4533, hsa-mir-548ac, hsa-mir-548i, hsa-mir-5585-3p, hsa-mir-6750-3p, hsa-mir-200c-3p, hsa-mir-1244, RERE(arginine-glutamic acid dipeptide repeats), HMG20B, MXD3, ARID4B, CBFB (core-binding factor subunit beta), TAF7 and CREM (cAMP response element modulator) might be the novel biomarker for HF.

The molecules HIM6, HIM10 obtained good binding score of more 5 to 6.999 with all proteins and the molecules HIM11, HIM12, HIM14, HTZ9, HTZ10 and HTZ12 obtained binding score above 5 and less than 9 with PDB protein code of 2HV7, 3VD8 and 6RUR respectively. The molecule HIM11 obtained highest binding score of 8.678 with 2HV7 and its interaction with amino acids are molecule HIM11 (Fig. 9) has obtained with a high binding score with PDB protein 2HV7, the interactions of molecule is the C6 side chin acyl carbonyl C=O formed hydrogen bond interaction with amino acid GLN-207 with bond length 1.92 A° and 3’ N-H group of imidazole ring formed hydrogen bond interaction with VAL-305 with bond length 2.36 A° respectively. It also formed other interactions of carbon hydrogen bond of -CH_3_ group of carboxylate at C6 with PRO-304 and amide-pi stacked and pi-pi stacked interaction of electrons of aromatic ring A with ALA-204 and ring C with HIS-155 and HIS-308. Molecule formed pi-alkyl interaction of ring B with PRO-304 and all interactions with amino acids and bond length are depicted by 3D and 2D figures (Fig.10 and Fig.11).

**Fig. 9.**
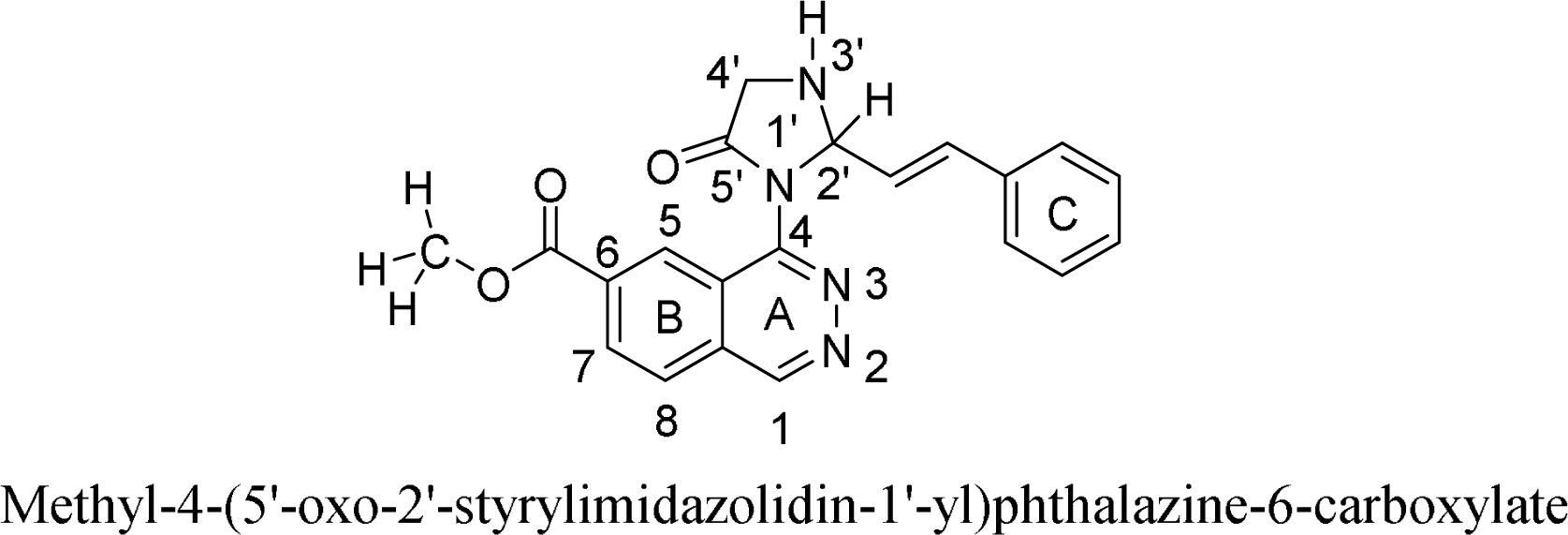
Structure of active designed molecule of HIM11

**Fig.10. 3D.**
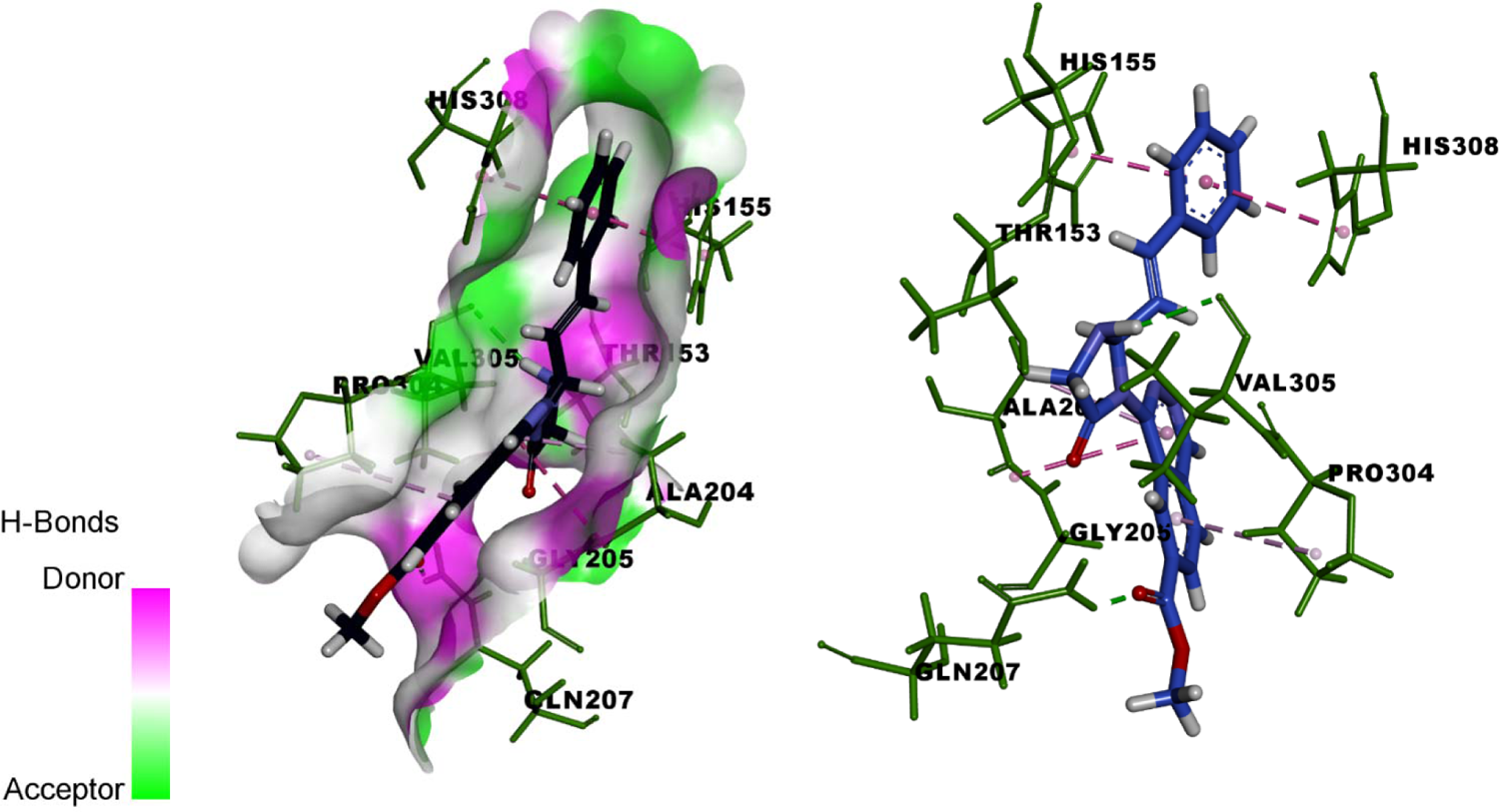
Binding of Molecule HIM11 with 2HV7

**Fig.11. 2D.**
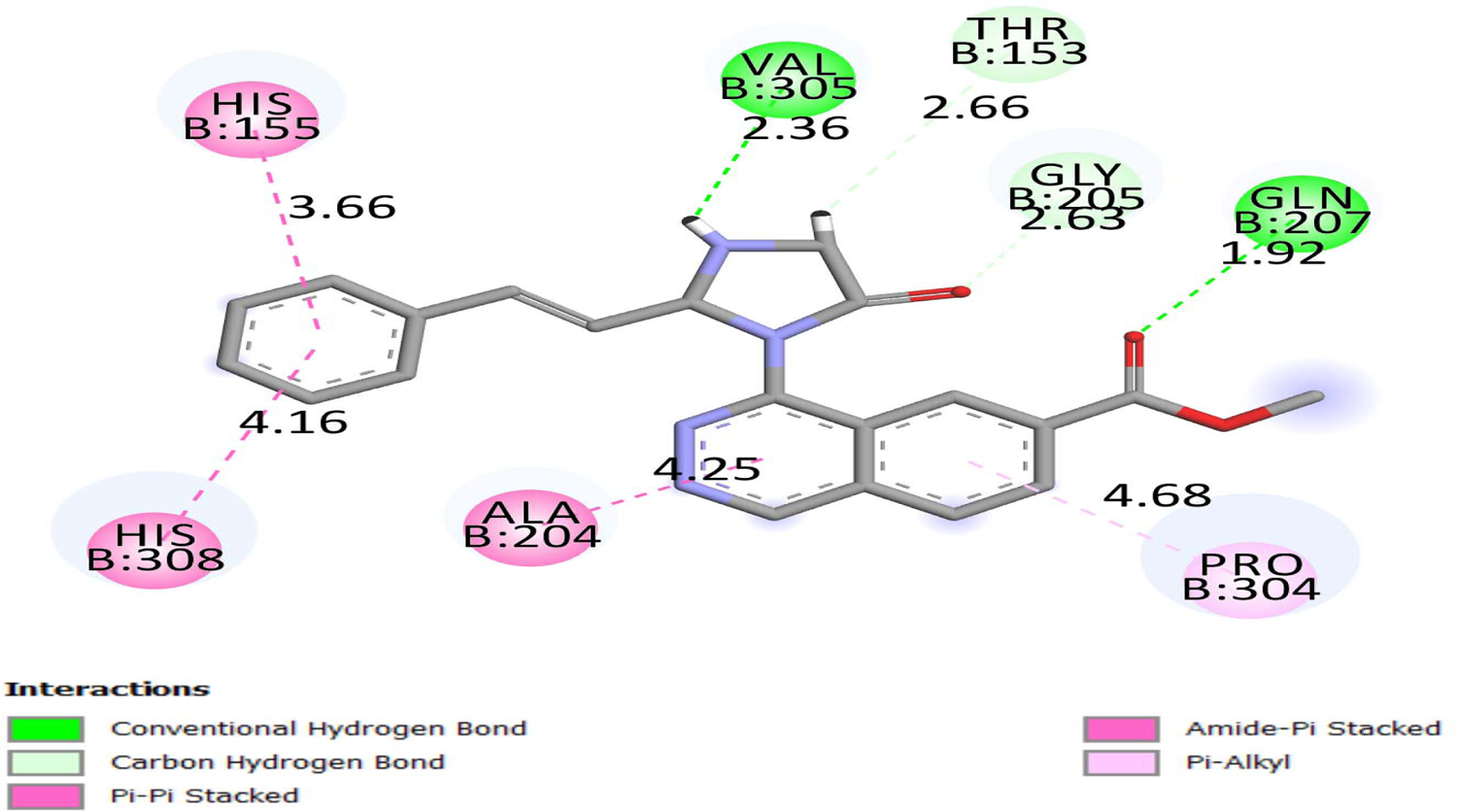
Binding of Molecule HIM11 with 2HV7

## Conclusions

The present study provided an extensive bioinformatics analysis of DEGs and revealed a series of targets and pathways, which may affect the progression of HF, for future investigation. And many of them, such as ESR1, PYHIN1, PPP2R2B, LCK, TP63, PCLAF, CFTR, TK1, ECT2 and FKBP5, might be key genes related to HF. The current investigation identified active molecules that might represent novel treatments for HF and may reduce the range of potential drugs for treating HF. These findings add to significant insights into the diagnosis and treatment of HF.

## Acknowledgement

I thank Michael Patrick Morley, Perelman School of Medicine at the University of Pennsylvania, Penn Cardiovascular Institute, Philadelphia, USA, very much, the author who deposited their profiling by high throughput sequencing dataset GSE141910, into the public GEO database.

## Ethical approval

This article does not contain any studies with human participants or animals performed by any of the authors.

## Informed consent

No informed consent because this study does not contain human or animals participants.

## Availability of data and materials

The datasets supporting the conclusions of this article are available in the GEO (Gene Expression Omnibus) (https://www.ncbi.nlm.nih.gov/geo/) repository. [(GSE141910) (https://www.ncbi.nlm.nih.gov/geo/query/acc.cgi?acc=GSE141910]

## Consent for publication

Not applicable.

## Author Contributions

B. V - Writing original draft, and review and editing

A. T - Formal analysis and validation

C. V - Software and investigation

## Authors

Basavaraj Vastrad ORCID ID: 0000-0003-2202-7637 Anandkumar Tengli ORCID ID: 0000-0001-8076-928X Chanabasayya Vastrad ORCID ID: 0000-0003-3615-4450

